# Set1/COMPASS repels heterochromatin invasion at euchromatic sites by disrupting Suv39/Clr4 activity and nucleosome stability

**DOI:** 10.1101/630970

**Authors:** R.A. Greenstein, Ramon R. Barrales, Nicholas A. Sanchez, Jordan E. Bisanz, Sigurd Braun, Bassem Al-Sady

## Abstract

Protection of euchromatin from invasion by gene-repressive heterochromatin is critical for cellular health and viability. In addition to constitutive loci such as pericentromeres and subtelomeres, heterochromatin can be found interspersed in gene-rich euchromatin, where it regulates gene expression pertinent to cell fate. While hetero- and euchromatin are globally poised for mutual antagonism, the mechanisms underlying precise spatial encoding of heterochromatin containment within euchromatic sites remain opaque. We investigated ectopic heterochromatin invasion by manipulating the fission yeast mating type locus boundary, using a single-cell spreading reporter system. We found that heterochromatin repulsion is locally encoded by Set1/COMPASS on certain actively transcribed genes and that this protective role is most prominent at heterochromatin islands, small domains interspersed in euchromatin that regulate cell fate specifiers. Interestingly, this effect can be gene orientation dependent. Sensitivity to invasion by heterochromatin, surprisingly, is not dependent on Set1 altering overall gene expression levels. At least two independent pathways direct this Set1 activity–inhibition of catalysis by Suv39/Clr4 and disruption of nucleosome stability. Taken together, these results describe a mechanism for spatial encoding of euchromatic signals that repel heterochromatin invasion.

## INTRODUCTION

Heterochromatin is a conserved nuclear ultrastructure [1], which enacts genome partitioning by repressing transcription and recombination at repetitive sequences and structural elements, as well as genetic information not pertaining to the specified cell fate. Once seeded at specific sequences [2–4], heterochromatin is subsequently propagated *in cis* over qualitatively distinct regions of the chromosome in a process termed spreading. Positional regulation of heterochromatin is key to determining and remembering cell fate decisions. Boundary regions often separate adjacent heterochromatin and euchromatin domains, reinforcing the distinct signals and functional environments on each side and countering the intrinsic propensity for heterochromatin to invade and silence genes. Major mechanisms of boundary formation fall into three broad classes: (1) recruitment of factors that directly antagonize the opposite state, for example by removal of state specific signals on chromatin [5–8]. (2) Promotion of the original state by either depositing or protecting such signals [9–12]. (3) Structural constraint via recruitment of DNA binding proteins that tether heterochromatin regions to the nuclear periphery [13–15]. Despite the varied modalities employed in boundary formation, containment is not absolute. This is evidenced by the observation that boundaries can be overcome by modest dosage changes in heterochromatin factors [13, 16], which leads to the silencing of genes critical to normal cellular function.

In addition to constitutive heterochromatin found at centromeres, telomeres, and other repetitive sequences, repressed domains also form at additional genomic locations in response to developmental and environmental signals [17–19]. These facultative heterochromatin domains are often embedded in euchromatic regions and silence developmental genes in a lineage specific manner [17]. Resulting from response to changing stimuli, the final extent of facultative domains can change over time, expanding to different degrees [17] and even contracting [20] in genomic space, though how this is achieved is not well understood. Facultative domain size may be tuned at the level of the heterochromatin spreading reaction [21] and/or the activities promoting its containment or disassembly. While little is known about the former, several models, beyond those known to operate at constitutive boundaries [19, 22], could be invoked to explain the latter.

How might euchromatin regulate heterochromatin spreading at facultative sites or respond to its expansion beyond constitutive domains, under conditions such as altered dosage regimes? One of the defining features of euchromatin is the presence of active genes. It is thought that transcription from active genes is incompatible with heterochromatin formation [23]. Multiple direct effects of transcription have been proposed to interfere with heterochromatin assembly. These include nucleosome turnover (eviction) by transcribing polymerase, formation of nucleosome-depleted regions at transcriptional units, or steric interference by transcription associated complexes [13, 24, 25]. Furthermore, we understand that unique molecular signatures characterize eu- and heterochromatin states and are critical to their formation. Heterochromatin is marked by methylation of histone 3 at lysine 9 or lysine 27 (H3K9me and H3K27me, respectively) and hypoacetylation of various lysine residues. In contrast, euchromatin features H3K4me, H3K36me and histone hyperacetylaton [22, 26]. Multiple studies have documented the apparent mutual exclusion of H3K9me- and H3K4me-marked regions [22, 26-28] and the requirement for removal of signals associated with the opposite state [6, 29]. While we are beginning to understand how this dichotomy is formed, it still remains unclear whether this is a cause or consequence of separating heterochromatin and euchromatin.

We aimed to investigate the role of euchromatic signals in regulating the extent of spreading in fission yeast, a well-characterized model system for the study of heterochromatin formation, which shares critical features with the processes found in metazoans. Fission yeast form constitutive heterochromatin marked by H3K9me at centromeres, telomeres, and the mating type (MAT) locus. Boundary formation occurs at peri-centromeric regions and the MAT locus via at least two mechanisms – tethering to the nuclear periphery through binding of TFIIIC proteins to *B-box* element sequences in boundary regions [30] as well as specific enrichment of a JmjC domain-containing protein, Epe1 [5, 7, 8, 31], which recruits additional downstream boundary effectors. In this system, facultative heterochromatin forms at developmentally regulated meiotic genes in regions surrounded by canonical euchromatin, which are partially dependent on Epe1 for containment [11, 19]. Utilizing the well-characterized MAT locus boundary as a model for euchromatic invasion, we found that active genes units could repel spreading and that this function depends on the H3K4 methylase complex Set1/COMPASS, independent of transcription. Set1 is the catalytic subunit of COMPASS and is responsible for mono-, di-, and tri-methylation of H3K4 *in vivo*. It is recruited by RNA polymerase and forms a characteristic pattern of H3K4 methylation states over genes, with H3K4me3 near the transcription start site (TSS) and H3K4me2 in the gene body (reviewed in [32]). We show that rather than acting as a global antagonist of spreading, like Epe1 or the histone acetyltransferase Mst2 [11], Set1 regulates spreading especially at gene-rich environments such as heterochromatin islands and that gene orientation can influence spreading outcomes. Set1 exerts its euchromatin protective function via at least two mechanisms: (1) the disruption of nucleosome stability and (2) catalytic inhibition of the sole fission yeast H3K9 methylase Suv39/Clr4, by the Set1 product H3K4me. This study provides a mechanism for the encoding of spatial cues within euchromatin that contain heterochromatin expansion.

## RESULTS

### Genes can function as a barrier to heterochromatin spreading

To investigate ectopic invasion of heterochromatin, we employed our previously described heterochromatin spreading sensor (HSS) [33, 34] in the euchromatic region proximal to the MAT Inverted Repeat Right (IR-R) boundary [35]. This HSS system contains two central components: (1) the spreading sensor, a monomeric Kusabira-Orange 2 fluorescent protein driven by the validated *ade6* promoter, hereafter referred to as “orange”, integrated 0.7kb outside IR-R, and (2) the control, a E2Crimson fluorescent protein driven by the same promoter, hereafter referred to as “red”, integrated at a constitutive euchromatic locus [33] (**Figure 1A**). We use flow cytometry to capture information from tens of thousands of single cells. With the HSS system, we normalize for cell-to-cell transcription and translation noise, allowing us to quantify heterochromatin-specific gene silencing at the “orange” reporter over the population.

**Figure 1:**
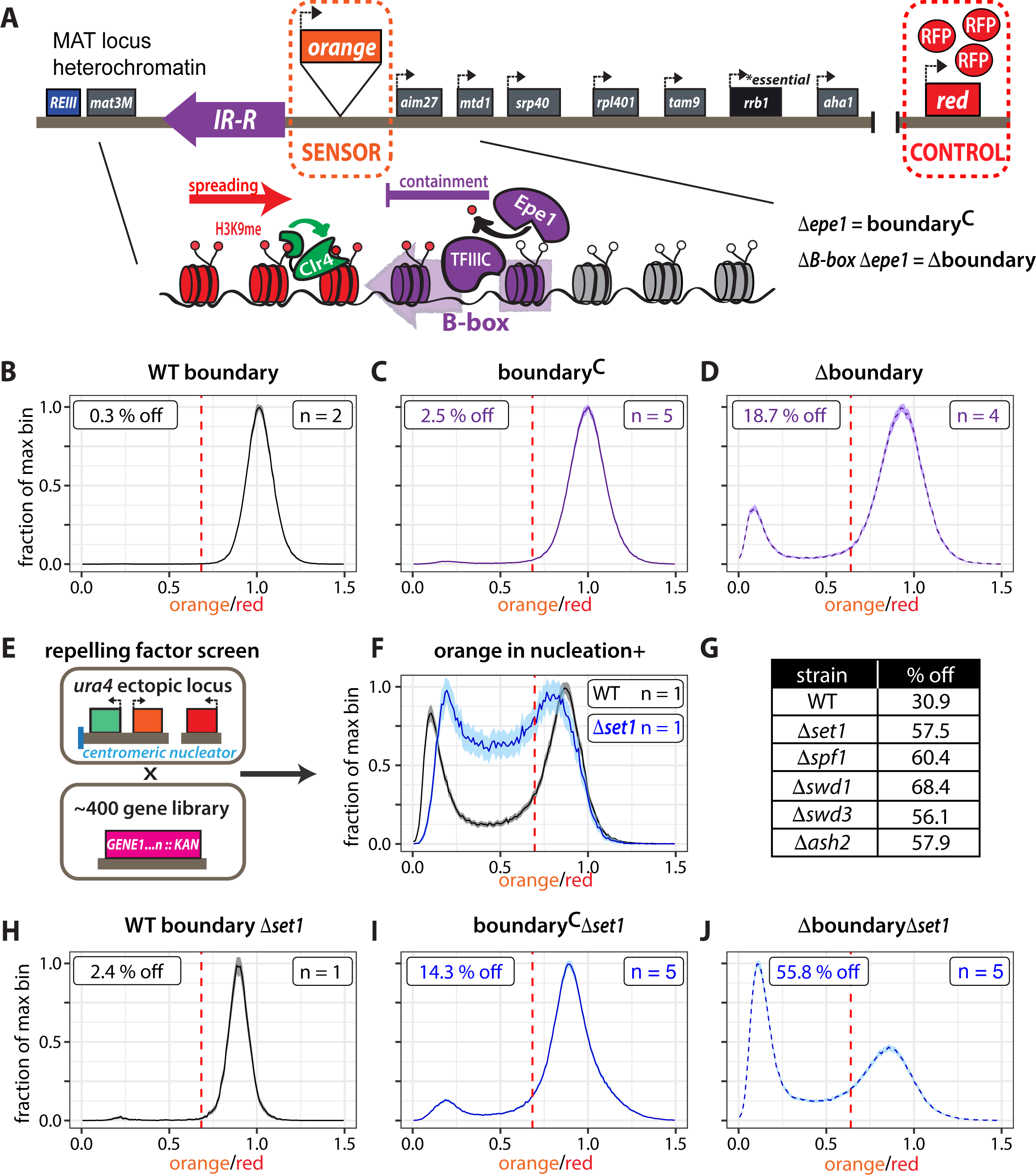
Genes repel heterochromatin across boundaries in a manner dependent on Set1/COMPASS. **(A)** An overview of the heterochromatin spreading sensor (HSS) outside the MAT locus IR-R boundary with transcriptional reporters encoding fluorescent proteins as sensor (“orange”) and control (“red”). IR-R (depicted as purple arrow) employs at least two independent pathways dependent on Epe1 and TFIIIC, respectively, to contain spreading of H3K9 methylation via Suv39/Clr4. IR-R function can be abrogated by deletion of *epe1* and removing the *B-box* binding sequences for TFIIIC. **(B)** Histogram of “orange” signal in a WT boundary background normalized to *Δclr4*. **(C)** Histogram of “orange” signal in boundary^C^ (*Δepe1*) background normalized to the corresponding WT (*epe1+*) strain **(D)** Histogram of normalized “orange” signal in Δboundary (*Δepe1 ΔB-box*) background as in C. **(E)** Illustration depicting genetic screen for modulators of gene-mediated heterochromatin repulsion. An HSS variant at the *ura4* locus was crossed to approximately 400 gene deletions. The resulting strains were analyzed by flow cytometry. **(F)** Histograms plotted as in C. of normalized “orange” signal in nucleation-gated cells in WT and *Δset1*. **(G)** Fraction of cells that experienced silencing at “orange”. Two thresholds were applied, a cutoff for nucleation at “green” and a cutoff for silencing at “orange”. Cells that met both criteria were counted as repressed. **(H)** Histogram of normalized “orange” signal in WT boundary*Δset1* background as in C. **(I)** Histogram of normalized “orange” signal in boundary^C^*Δset1* background as in C. **(J)** Histogram of normalized “orange” signal in WT Δboundary*Δset1* background as in C. All 1D histograms are plotted as the mean +/-3SD of 300 bootstrap iterations for combined data from the indicated number of biological isolates (n). Signal is normalized to the median signal from a *Δclr4* or corresponding WT (*epe1+*) strain control to represent the maximum fluorescence in the absence of heterochromatin (x=1). A threshold for silencing (red line) represents the mean signal of the WT strain less 2SD with the exception of (F) where the threshold for silencing in nucleation+ cells was determined as mean less 1SD of the “orange” signal from the *Δclr4* strain. The faction of cells below this cutoff was calculated (%off).

We first examined the normalized (legend to Figure 1 and [33]) orange fluorescence of a strain with a WT boundary (*epe1+, B-box+*) and detect no silencing in the population distribution (**Figure 1B**). We define a threshold for silencing as the mean of the appropriate WT (*epe1+*) strain less two standard deviations. We next compromised one or both of the pathways required for containment of spreading at IR-R [7, 8, 36] and assessed the effect on “orange” silencing. Consistent with previous results [36], very little silencing is detected in *Δepe1* isolates harboring a partially compromised boundary (referred to hereafter as boundary^C^) (**Figure 1C**). In a fully compromised boundary absent both *epe1* and the 5 *B-box* sequence elements contained within IR-R [30] (referred to hereafter as Δboundary), we detected increased silencing (**Figure 1D**). Yet, even in the Δboundary background, greater than 80% of cells in the population fully express “orange”. Given this result, and the observation that H3K9me2 spreading declines sharply over endogenous IR-R bordering genes [36], we wondered whether activities acting at the reporter and potentially endogenous gene(s) themselves repel spreading.

### Set1/COMPASS regulates genic protection from heterochromatin spreading

In order to identify potential factors that regulate gene-mediated repulsion of heterochromatin spreading, we designed a genetic screen to query the effect of gene deletions on silencing measured via our reporters. To do this, we moved the HSS to the euchromatic *ura4* locus (**Figure 1E**) downstream of a previously described RNAi-based heterochromatin nucleator [37]. We have demonstrated that this construct can generate spreading up to 8kb downstream [33] without the need to remove any boundary factors, thus avoiding confounding global effects on growth in the screen. At this locus, a reporter cassette encoding “green”, a third fluorescent protein driven by *ade6p*, is integrated 1kb downstream of the RNAi element while the “orange” cassette is integrated 3kb downstream from “green” in the same orientation with respect to spreading as at IR-R. While nucleation at this locus is not as robust compared to endogenous heterochromatin domains, we can apply a computational gate to isolate successfully nucleated cells (greenOFF, described in [33]) and assess their spreading state. In the WT background, the nucleation gated “orange” signal in this strain resembles the behavior seen in the Δboundary IR-R HSS strain (compare **Figure 1F**, black line and **Figure 1D**), exhibiting both gene silencing and fully expressed states.

We crossed this *ura4*-HSS background strain to a curated ∼400 gene subset of the *S. pombe* deletion library enriched for nuclear factors (**Figure 1E**) and measured reporter fluorescence from the resultant strains via flow cytometry. For each strain, we plotted a 2D histogram of red-normalized orange versus green fluorescence (**Supplemental Figure 1**) and calculated the fraction of cells that experienced silencing at “orange”. Silencing in this context is defined as the fraction of all cells that met both the greenOFF criteria for nucleation (blue line) and had orange signal below the mean less 1 standard deviation of the matched *Δclr4* strain (red line). We use 1 standard deviation as a cutoff for the screen given the relatively broader distribution of the “red” control in this background. The majority of this screen will be described elsewhere.

Upon analysis of this dataset, we noticed 5 genes whose absence had the same characteristic effect of increased silencing at the spreading reporter – *ash2, swd1, swd3, spf1,* and *set1* (**Figure 1F, Figure 1G, Supplemental Figure 1**). These are five members of the Set1/COMPASS complex, which catalyzes H3K4me and deposits H3K4me3 at active gene promoters [38–41]. Of the remaining members, *Δswd2* did not grow and *Δsdc1* was not in the screen, while *Δshg1* showed no phenotype, consistent with other studies, which denote it marginally associated with the complex [40]. All five gene deletions were validated by independent knockout in the parental reporter background.

Given this result, we sought to test whether the removal of Set1C might have a similar effect at the boundary proximal locus. While there was not a major effect of *Δset1* on reporter strains with a WT boundary (**Figure 1H**), both boundary^C^ (**Figure 1I**) and Δboundary (**Figure 1J**) proximal reporters experienced a significant increase in silencing in *Δset1*, supporting the hypothesis that the reporter gene itself was capable of blocking spreading and that this protective function depends on Set1/COMPASS.

### Set1 contributes to containment of heterochromatin spreading globally

In order to probe the effect of *Δset1* on euchromatic invasion at heterochromatic sites genome wide, we performed Chromatin Immunoprecipitation followed by Next Generation Sequencing (ChIP-Seq) with antibodies against H3K4me3 and H3K9me2 in WT, *Δepe1*, and *Δepe1Δset1* strains that contained no reporters (**Figure 2A**). We did not perform H3K4me3 ChIP-Seq for *Δset1* isolates due to the absence of H3K4me, which we validated by ChIP-qPCR (**Supplemental Figure 2E**). Signal tracks for each genotype are plotted as mean and 95% confidence interval of 2-4 replicates.

**Figure 2:**
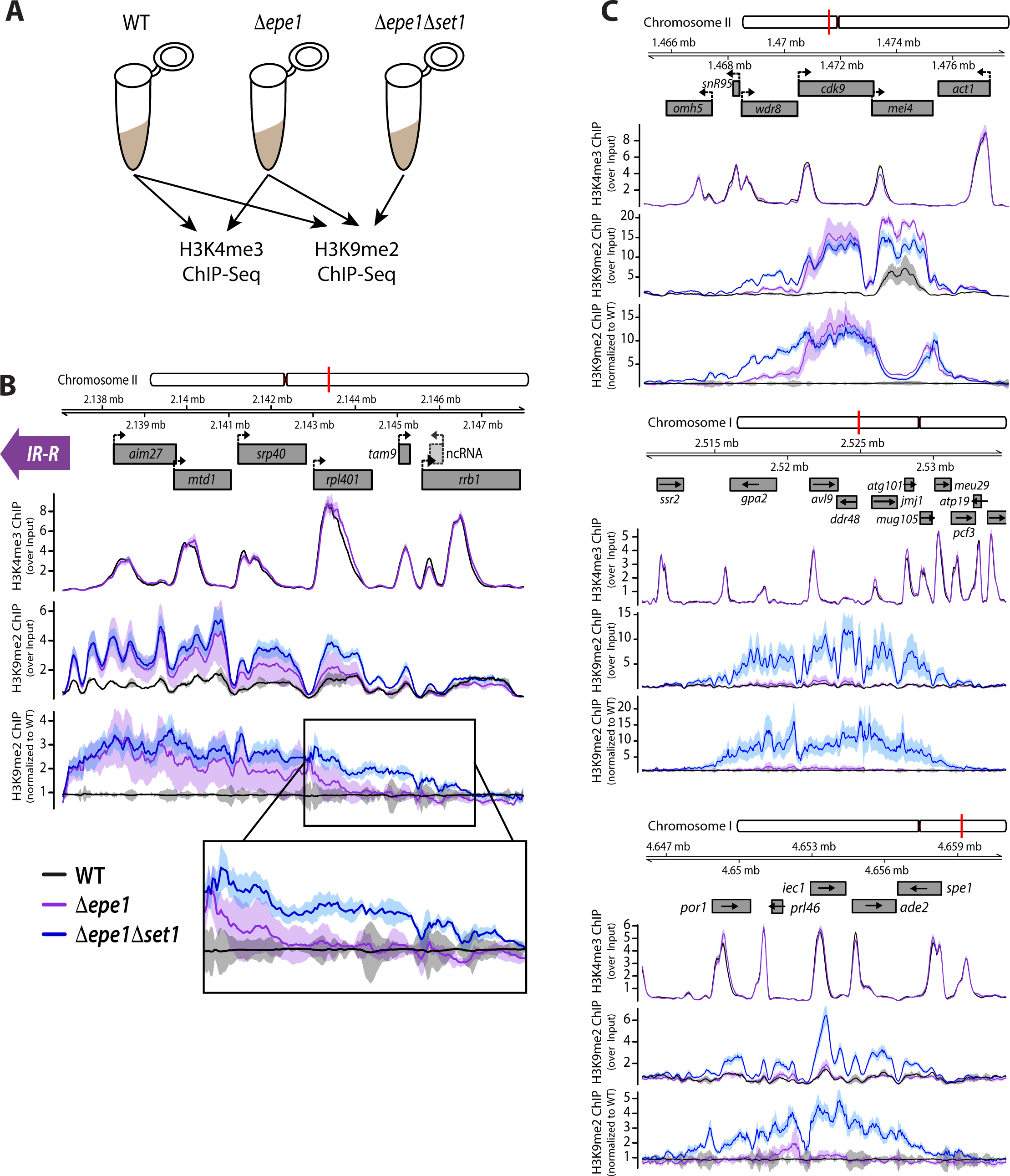
Set1 regulates H3K9me spreading at gene-rich euchromatic loci. **(A)** Overview of ChIP-Seq experiment. **(B)** ChIP-Seq signal tracks and gene annotations for the IR-R proximal region. **(C)** Signal tracks and annotations for three regions previously identified as heterochromatin islands. Each sample including the input was normalized as signal per million reads and then ChIP samples for H3K4me3 and H3K9me2 were normalized to Input (top and middle tracks). H3K9me2 ChIP-Seq datasets were independently normalized to signal from a sample containing merged data from both WT isolates (bottom track). Signal tracks are plotted as the average value per 25bp bin. Tracks are represented as mean (line) and 95% confidence interval (shaded region) per genotype (WT n=2, *Δepe1* n=3, *Δepe1Δset1* n=4; each n represents a single colony deriving from an original knockout for each genotype).

Given that our above results show *set1*-dependent heterochromatin repulsion by our reporter gene, we asked whether the removal of *set1* would affect spreading beyond IR-R (**Figure 2B**). While no enrichment of H3K9me2 was detected beyond IR-R in WT (black line), both *Δepe1* (purple line), and *Δepe1Δset1* (blue line) display similar and significant enrichment for H3K9me2 immediately next to IR-R, as seen by their closely superimposed means and confidence intervals. As distance increases from IR-R, the traces begin to separate visibly, with H3K9me2 signal from *Δepe1Δset1* strains significantly exceeding that from *Δepe1* and WT. This separation is most evident over the open reading frame of *rpl401* (**Figure 2B**, inset). Interestingly, this gene is also highly enriched for H3K4me3. The signal in *Δepe1Δset1* remains significantly above WT levels up until it reaches the essential gene *rrb1*. Given this result, we conclude that *set1* contributes to the containment of spreading into the euchromatic region outside of IR-R in the case of boundary failure.

Encouraged by the result at IR-R, we next examined other constitutive heterochromatin loci for *set1*-mediated spreading effects. Broadly, *Δset1* did not significantly increase the extent of spreading already evident in *Δepe1* at such loci. Increased spreading was detected in *Δepe1Δset1* beyond the boundaries of pericentromeric heterochromatin on chromosome II and III (**Supplemental Figure 2B**), while at the right subtelomere I and at the pericentromere of chromosome I spreading was in fact reduced in *Δepe1Δset1* relative to *Δepe1* (**Supplemental Figure 2C**).

Given the major role of Set1/COMPASS at genes and the enrichment of H3K4me in canonical euchromatin [26], we wondered if *set1* might regulate spreading at facultative heterochromatin sites, islands of H3K9me embedded in gene-rich euchromatin [11, 19]. We found that even the core of such islands maintain TSS-proximal H3K4me despite being marked by low to intermediate levels of H3K9me2 **(Figure 2C)**. This is consistent with a previous report that measured H3K4me2 by ChIP-chip [19]. Our analysis determined a similar overall number of previously described heterochromatin islands and novel ectopic H3K9me2 peaks (sites where the WT stains shows no significant H3K9me2 enrichment) between *Δepe1* and *Δepe1Δset1* mutants (**Supplemental Figure 2A**). However, we found that heterochromatin spreading is exacerbated significantly in *Δepe1Δset1* compared to *Δepe1* at several sites: Specifically, we detected increased H3K9me2 spreading at 2 islands (**Figure 2C**, centered on *mei4* and *iec1*), and an additional 3 ectopic sites **(Figure 2C** and data not shown). Further, we identified one site that was formed exclusively in the *Δepe1Δset1* (**Supplemental Figure 2D**), and not *Δepe1*. These results describe a role for *set1* in spreading containment at gene-rich euchromatin with prominent H3K4me3 peaks.

### Set1 confers barrier activity to MAT-adjacent gene promoters but does not regulate their steady state transcription

We next wanted to test whether endogenous boundary proximal genes could function as barriers to ectopic heterochromatin invasion and if so, whether *set1* mediated their ability to repel heterochromatic silencing. To address this question, we chose to test two genes from the IR-R adjacent region that were enriched for H3K9me2 in the absence of *epe1*: (1) *rpl401,* the gene at which the *Δepe1Δset1* double mutant first significantly exceeds *Δepe1* alone, and (2) *mtd1*, the locus where we first start to detect visible, although not statistically significant, separation in the H3K9me2 tracks between the genotypes (**Figure 2B**).

We first modified the original *ade6p*-HSS to express “orange” from the *rpl401* promoter at the same locus (**Figure 3A**). The *rpl401* gene promoter effectively repels spreading in the context of a compromised (boundary^C^, **Figure 3B**) or fully abrogated (Δboundary, **Figure 3C**) IR-R boundary. This may not be too surprising given that *rpl401* is a very highly transcribed gene [42]. However, the removal of *set1* (*Δset1*) resulted in complete *rpl401* repression in a Δboundary context (**Figure 3C**). This indicates that both *ade6p* and *rpl401p* can form a spreading barrier that is highly sensitive to the presence of Set1. To examine spreading at the site of an endogenous gene, instead of at an inserted reporter, we replaced the *mtd1* open reading frame with “orange” to generate an *mtd1p-*HSS (**Figure 3D**), which is located 2.5kb from the edge of IR-R. Just like *ade6p-*HSS and *rpl501p-*HSS at the IR-R proximal locus, the *mtd1p-*HSS also displays genic barrier function that is *set1-*dependent (**Figure 3E**). However, unlike *ade6p-*HSS and *rpl401p-*HSS, the *mtd1p-*HSS is completely repressed (**Supplemental Figure 3A**) when IR-R is fully abrogated. This suggests that while all the gene promoters we examined are sensitive to *set1* for barrier function, some constitute only weak barriers, like *mtd1*.

**Figure 3:**
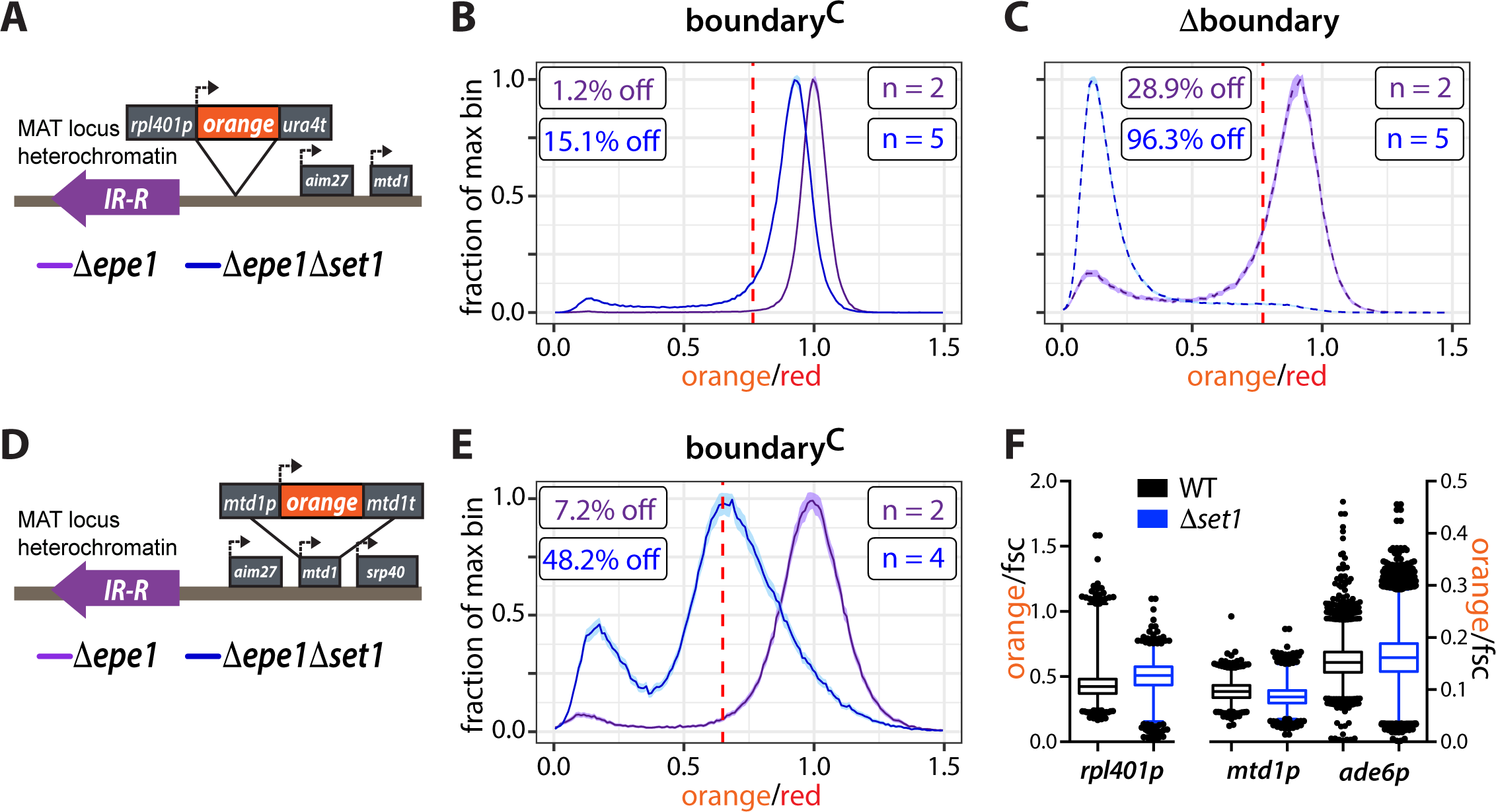
Set1 regulates genic heterochromatin barrier function of MAT-adjacent gene promoters but does not regulate their transcription. **(A)** Overview of the *rpl401p*-HSS. **(B)** Histogram plots as in Figure 1 of normalized “orange” signal from *set1*+ (purple) and *Δset1* (blue) *rpl401p*-HSS boundary^C^ isolates. **(C)** Histogram plots of normalized “orange” signal from *rpl401p*-HSS Δboundary isolates. **(D)** Overview of the *mtd1p*-HSS. **(E)** Histogram plots of normalized “orange” signal from *mtd1p*-HSS boundary^C^ isolates. **(F)** Box and whisker plots of “orange” signal normalized to forward scatter (fsc) for *rpl401p-, mtd1p-,* and *ade6p-*HSS in *set1*+ (black) and *Δset1* (blue) backgrounds. 1%-99% of the data is included within the whiskers. Outliers are plotted as individual points.

Previous reports have described both transcription activating and repressive roles for Set1/COMPASS [43–46]. To directly test whether *set1* might confer genic-protection from heterochromatin spreading by altering gene transcript levels, we examined the “orange” signal expressed from *rpl401p, mtd1p,* and *ade6p* in a *set1+* or *Δset1* backgrounds in a WT boundary context (**Figure 3F**). “orange” signal was normalized to forward scatter (fsc) as a proxy for cell volume, bypassing any confounding effect *Δset1* might have on our *ade6p*-driven “red” control. We did not detect any major decrease in “orange” in *Δset1* isolates (**Figure 3F**). We confirmed this result by RT-qPCR analysis, where we normalized *ade6p* “orange”, *mtd1*, and *rpl401* transcripts to an *act1* control (**Supplemental Figure 3C**). In the normalization, we adjusted for the *Δset1* effect on this control (**Supplemental Figure 3B**, methods). Together, these results argue against the hypothesis that Set1 regulates the mean level of RNAPII mediated transcription at these genes. Thus, it is unlikely that altered frequency of RNAPII passage through local genic chromatin causes the *Δset1-*dependent phenotype in reporter silencing.

### H3K4me abrogates spreading by direct interference with Suv39/Clr4 catalysis

SetC is the only H3K4 methylase in fission yeast [38] and H3K4me and H3K9me appear mutually exclusive [26]. Hence, we sought to test whether the biochemical basis for the *set1*-mediated genic barrier phenomenon could be caused by H3K4me-mediated inhibition of the heterochromatin spreading reaction. This could potentially occur via two mechanisms – either by directly impacting catalysis of H3K9 methylation by Suv39/Clr4 or by disrupting the ‘read-write’ positive feedback characteristic of histone methyl transferases. The spreading feedback mechanism is mediated by the binding of Suv39/Clr4 enzyme to its own product via the chromodomain (CD), which stimulates the catalysis of H3K9 methylation on proximal nucleosomes via the SET domain [47–49].

In support of the feedback model, it was previously shown that acetylation (ac) of H3K4 inhibits the binding of the Clr4-CD to H3K9me3 tail peptides [50]. This suggests steric hindrance to binding might be caused by additional posttranslational modifications at the H3K4 residue. We therefore tested whether H3K4me3, which is primarily present around the TSS, could regulate product recognition. We purified the Clr4-CD (**Supplemental Figure 4A**) and performed fluorescence polarization with modified histone tail peptides. We found that the Clr4-CD has a similar binding affinity for H3K9me3 and H3K4me3K9me3 tail peptides, with a K_d_ of 1.97 +/-0.05μM and 1.85 +/-0.16μM respectively (**Figure 4A**). Thus, the presence of H3K4me3, unlike H3K4ac, does not disrupt the ‘read-write’ feedback mechanism.

**Figure 4:**
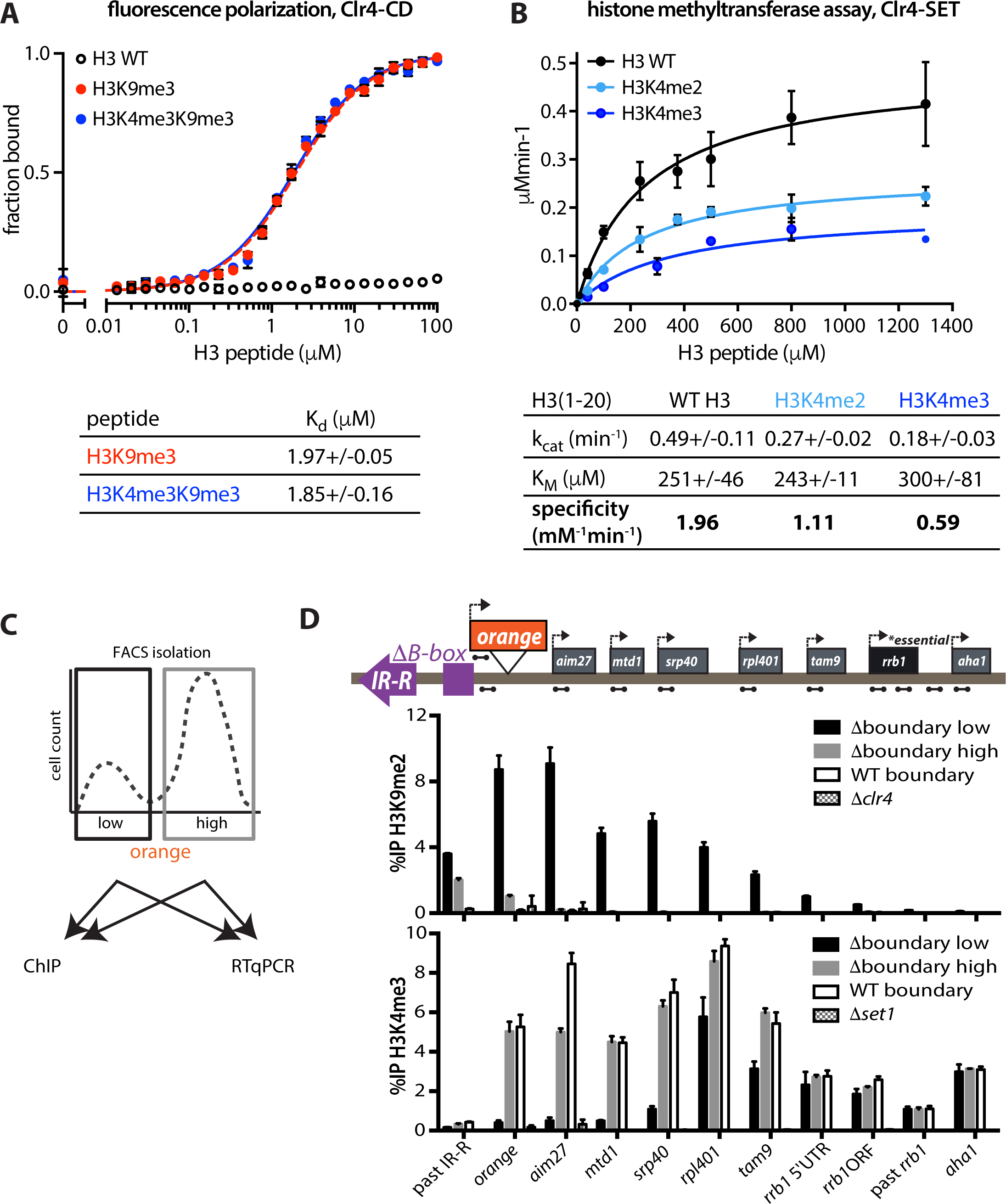
Gene-protective activity is rooted in catalytic inhibition of Suv39/Clr4 by H3K4me2/3. **(A)** Fluorescence polarization for the Suv39/Clr4 Chromodomain and fluoresceinated H3(1-20) peptides with modifications as indicated. **(B)** Histone methyltransferase assay with Clr4-SET and H3(1-20) peptides with modifications as indicated. Error bars in A. and B. represent 1SD from three replicate experiments. Calculated kinetic parameters are indicated in the table. Values represent mean and 1SD of three independent curve fits. **(C)** Cartoon overview depicting FACS isolation of “low” and “high” Δboundary 5′ *ade6p-*“orange” cells followed by ChIP and RT-qPCR. **(D)** ChIP-qPCR data for FACS sorted cells - H3K9me2 (top) and H3K4me3 (bottom). Amplicons for each qPCR are depicted as dumbbells on cartoon locus. Error bars represent 1SD from three technical replicate ChIPs.

A previous study indicated no obvious effect of H3K4me2 on Suv39/Crl4 activity [51], yet a number of other studies document a range of effects of H3K4me2 or H3K4me3 on H3K9 methyl transferases, although these results are conflicting in specific cases [52–54]. Most of these studies carried out end point analysis. To definitively determine any effect of H3K4me2 or H3K4me3 may have on Suv39/Clr4 catalysis, we performed multiple turnover Michaelis-Menten kinetic analysis using N-terminal truncation of Clr4 comprising residues 192-490 [55, 56] which includes the catalytic SET domain (**Figure 4B** and **Supplemental Figure 4B**). We determined k_cat_ and K_M_ and specificity constant (k_cat_/K_M_) values (**Figure 4B**) and importantly found that H3K4me3 and H3K4me2 reduce Clr4’s k_cat_/K_M_ by 3.3 times and 1.8 times, respectively, relative to a H3K4me0 peptide. This derives mostly from an adverse effect on Suv39/Clr4’s k_cat_ rather than on the K_M_ (see **Figure 4B**). In conclusion, these results demonstrate that Suv39/Clr4 catalysis, but not its product recognition, is inhibited by the presence of H3K4me3, to and a milder extent, by H3K4me2.

Given this result, we wanted to test whether H3K4me accumulation at barrier genes protects the euchromatic state of transcriptional units downstream. Using Fluorescence Assisted Cell Sorting (FACS), we isolated both repressed (“low”) and expressed (“high”) populations of 5′ *ade6p*-HSS Δboundary cells (**Figure 4C**) and assessed their chromatin via ChIP and transcriptional state via RT-qPCR (**Figure 4D, Supplemental Figure 4C**). While both populations evidenced H3K9me2 accumulation upstream of the reporter, H3K9me2 signal cannot be detected at any point beyond “orange” in the “high” cells (grey bars). This immediate drop coincides with the *ade6p* H3K4me3 peak, and H3K4me3 remains significantly enriched at the downstream gene promoters, comparable to WT levels. Consistent with this H3K4me3 distribution, transcription levels are similar to the no heterochromatin (*Δclr4*) state. This result, in conjunction with our above findings (**Figure 1I,J** and **Figure 4B**) suggest that H3K4me3 accumulation at *ade6p* protects downstream transcriptional units. On the other hand, the “low” population (black bars) displays high levels of H3K9me2 at and beyond “orange”, while H3K4me3 is severely reduced (**Figure 4D**). H3K9me2 levels eventually decline towards the essential *rrb1* gene, concomitant with a rise in H3K4me3 enrichment. The difference between the H3K4me3 signal in the “low” and “high” populations thus eventually decreases with distance. In cells where *ade6p* localized H3K4me3 is overcome, downstream transcriptional units therefore appear to succumb to repressive H3K9me2.

We wondered whether mutations to the Suv39/Clr4 SET domain that reduced catalytic inhibition by H3K4me2/3 could overcome genic barriers *in vivo*. However, aspartate to alanine mutations homologous to those made in SetDB1 [54], resulted in a complete loss of silencing (data not shown), suggesting these mutations abrogated Suv39/Clr4 SET domain function. Nevertheless, the above results support the model that a gene can serve as a boundary to heterochromatin spreading to preserve the euchromatic state beyond it, and that H3K4 methylation enacts at least part of this barrier activity by direct inhibition of Suv39/Clr4 described above.

### Gene orientation-dependent and -independent barrier function

The TSS-proximal enrichment for H3K4me3 [41, 57-59] would predict that genes might create a stronger barrier to spreading when oriented with their 5′ end facing the expanding edge of an invading heterochromatin domain. We tested this hypothesis directly by comparing variations of our HSS with reporter cassettes integrated adjacent to IR-R oriented with either their promoter (5′ end) or terminator (3′ end) closer to heterochromatin. The original *ade6p*-, *rpl401p-,* and *mtd1p-*HSS constructs are in the 5′ proximal orientation with respect to IR-R. We generated 3′ proximal versions of these reporters (**Supplemental Figure 5A-J**) and assessed the fraction of cells that displayed silencing.

In agreement with this prediction, 3′ proximal reporters driven by the *ade6* promoter were significantly more repressed than 5′ proximal reporters in both boundary contexts (**Figure 5A**, purple bars and table). 3′ proximal *ade6p*-HSS in the Δboundary background was completely repressed (**Supplemental Figure 5C)** in contrast to 18.7% in the 5′ orientation (**Figure 1D**). However, this 5′ protection bias was not evident for the *rpl401* promoter: 5′ and 3′ proximal *rpl401p*-HSS reporters were equally repressed in boundary^C^ and the 5′ proximal reporter was marginally more repressed than the 3′ proximal version in the Δboundary context (**Figure 5B**, purple bars and table). Reporters transcribed from *mtd1p* were similarly repressed in both orientations (**Supplemental Figure 5G-I, Figure 3E, Supplemental Figure 3A**). Nevertheless, gene silencing in all promoters and orientations was significantly increased when *set1* was deleted (**Figure 5A,B** blue bars, **Supplement Figure 5A-I**).

**Figure 5:**
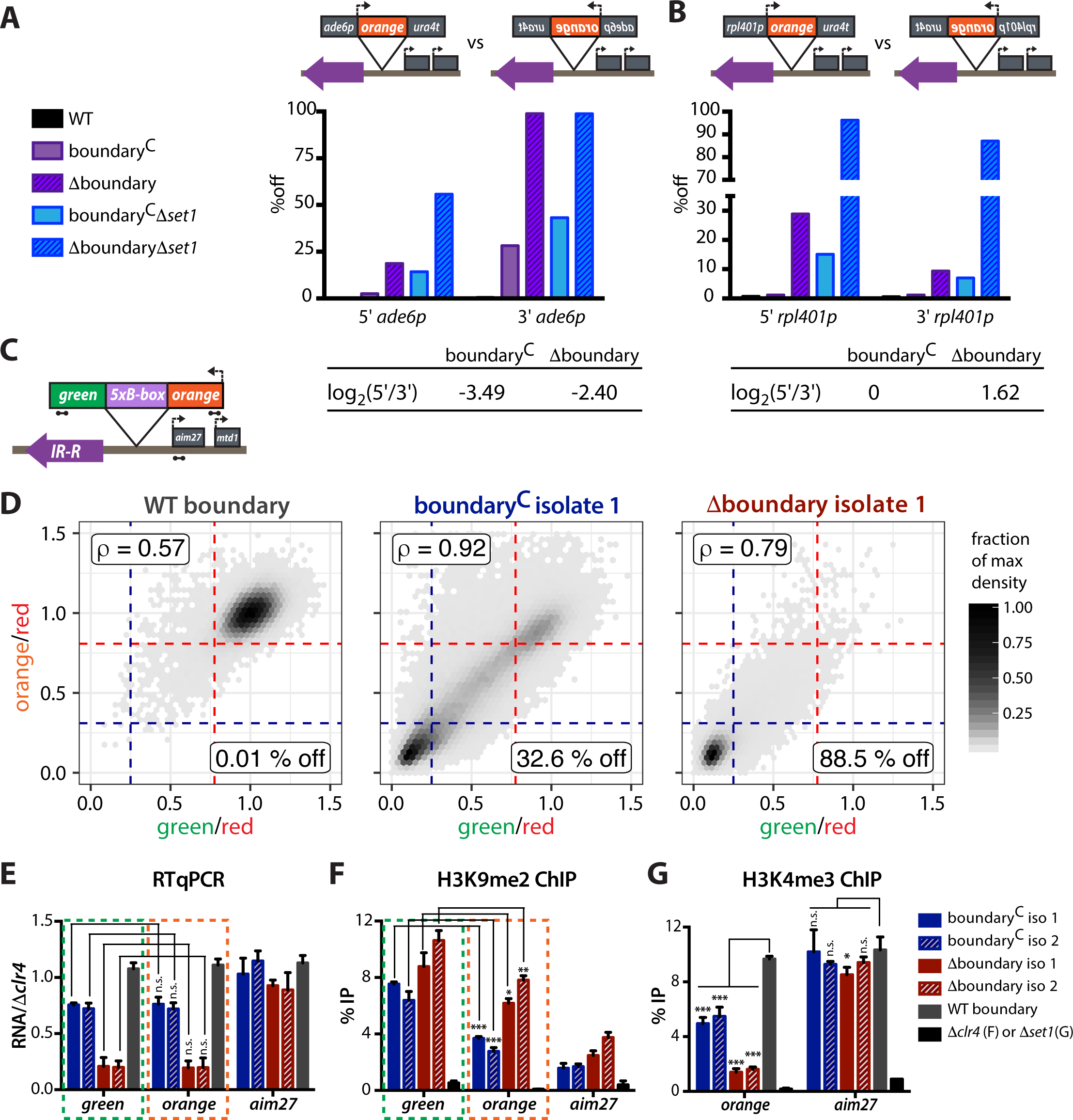
A 5′ protection effect is promoter specific and results from H3K9me spreading into the gene body. **(A)** Bar plots representing percentage off for *ade6p*-driven “orange” reporters in either the 5′ or 3′ orientation with respect to IR-R (cartoon). Boundary and *set1* genotypes are as indicated. **(B)** Bar plots as in A. representing percentage off for *rpl401p*-driven “orange” reporters in either the 5′ or 3′ orientation with respect to IR-R (cartoon). For A. and B. the orientation score, calculated as the log_2_ ratio of “%off” values for the 5′ orientation over 3′ orientation per boundary context, is denoted in a table below the respective plot. **(C)** Locus cartoon for 3′ *ade6p* construct which expresses “orange” and “green” ORFs joined by an in-frame linker containing 5 *B-box* sequences. **(D)** 2D density hexbin plots of normalized green and orange signal for WT, boundary^C^, and Δboundary isolates. All plots are normalized to the median signal from the WT boundary strain. Lines represent “on” (red) and “off” (blue) cutoffs. “On” is defined as mean of WT less 2SD while “off” is defined as the mean plus 2SD of a “red”-only control strain. Pearson correlation (ρ) for normalized “green” and “orange” and percentage of values less than the “off” cutoff for both colors is annotated. **(E)** RT-qPCR data from two boundary^C^, two Δboundary, and one WT isolate. **(F)** H3K9me2 ChIP-qPCR data from two boundary^C^ and two Δboundary isolates with *Δclr4* negative control. **(G)** H3K4me3 ChIP-qPCR data from two boundary^C^, two Δboundary and one WT isolate with *Δset1* negative control. Amplicons for each qPCR are depicted as dumbbells on cartoon locus. Error bars represent 1SD of three technical replicate ChIPs. n.s. represents P>0.05, * represents P<0.05, ** represents P<0.01, *** represents P<0.001 (t-test).

How can a gene’s barrier activity be 5′ orientation-sensitive, or -insensitive, and preserve its dependence on Set1, which is known to bias H3K4me3 deposition near the TSS [32, 41]? In what follows, we provide mechanisms that can account for *set1-*dependent gene barrier activity that is either orientation-sensitive or - insensitive.

### Invasion into the gene 3′ is sufficient for silencing and relief of inhibitory H3K4me3

First, we describe a mechanism underlying 5′ orientation dependence. We hypothesized that for gene barrier activity to be ineffective in the 3′ orientation, heterochromatin must be able to at least partially invade the gene from the 3′ end, which is depleted for H3K4me3 [32]. This would be consistent with our previous results that showed that a 3′ oriented reporter can experience intermediate repression, which correlated with intermediate levels of RNA, H3K4me3, and H3K9me2 [33]. Partial invasion may then lead to transcriptional silencing, which would inherently down-regulate H3K4me. This is because H3K4me deposition depends on signals from active RNA polymerase (reviewed in [32]). Removing the kinetic inhibition of H3K9 methylation then reinforces the repressed state. This notion is supported by our experiments at the *ura4* locus, where *ade6p-* driven “green” is oriented 3′ proximal to heterochromatin and spreading can overtake the gene unit and continue up to 7kb downstream (**Figure 1F** and [33]).

To test this hypothesis, we built a variant of the 3′*ade6p*-HSS reporter construct that would permit spreading to proceed into the gene unit but prevent it from reaching the promoter, mimicking an intermediate step in the process of gene invasion. To achieve this, we fused the “orange” and “green” coding sequences by an in-frame linker containing 5 B-box elements (**Figure 5C**). Previous reports demonstrated 3 B-box elements were sufficient to confer synthetic boundary activity [13]. Despite varying amounts of repression, signal from “green” and “orange” in WT, boundary^C^, and Δboundary contexts (**Figure 5D, Supplemental Figure 5B**), as well as their RNA levels (**Figure 5E**), were well correlated in each isolate. cDNA synthesis for RT-qPCR was performed with random hexamers to permit detection of any partial transcripts, however RNA levels for both XFP’s were equivalent in each strain. This indicates that the entire transcriptional unit is uniformly regulated, despite presence of the synthetic B-box boundary midway through the tandem gene unit. We next assessed the chromatin state at “green” and “orange” by ChIP. H3K9me2 is significantly reduced at “orange” compared to “green” across all isolates from both boundary^C^ and Δboundary contexts (**Figure 5F**) validating that the 5x B-box sequence was functioning as a synthetic boundary. Surprisingly, H3K4me3 ChIP revealed that boundary^C^ and Δboundary had significantly reduced methylation levels compared to WT at the “orange” TSS (**Figure 5G**). These results demonstrate that invasion of a gene from the 3′ end can reduce both inhibitory H3K4me levels and transcription, despite not reaching the gene promoter. This suggests a mechanism for silencing of 3′ genes in which, as the spreading machine invades a gene unit, heterochromatin formation over part of the gene body is sufficient to down-regulate transcription and H3K4me3, relieving inhibition of Suv39/Clr4 to facilitate invasion. Presumably this mechanism would not be able to operate in the 5′ orientation since H3K4me3 would be encountered first, likely preceding effective transcriptional downregulation.

### Set1 destabilizes nucleosomes globally, providing a gene orientation-insensitive genic barrier mechanism

Next, we hypothesized that the lack of orientation sensitivity in the barrier activity of *rpl401* and *mtd1* as well as lack of orientation bias in genes located proximal to constitute boundaries (**Supplemental Figure 6A**), could be explained by Set1/COMPASS-dependent heterochromatin antagonizing pathways beyond direct H3K4me inhibition. As transcription of boundary proximal genes is not reduced in *Δset1* (**Figure 3F, Supplemental Figure 3C**), we asked whether the deletion of *set1* affected two additional parameters known to interfere with heterochromatin formation – nucleosome occupancy [25] and histone acetylation [24]. We first assessed nucleosome occupancy in *set1+* and *Δset1* strains by H3 ChIP in log phase cultures (**Figure 6A**). This experiment revealed that *Δset1* led to an increase in nucleosome occupancy at euchromatic sites, while heterochromatin targets were unaffected. To exclude any possible effects due to passage through the cell cycle, we performed a similar analysis in G2 stalled cells via anti-HA ChIP in strains expressing a C-terminal HA fusion of one of the three H3 genes, which confirmed this result (**Supplemental Figure 6B**).

**Figure 6:**
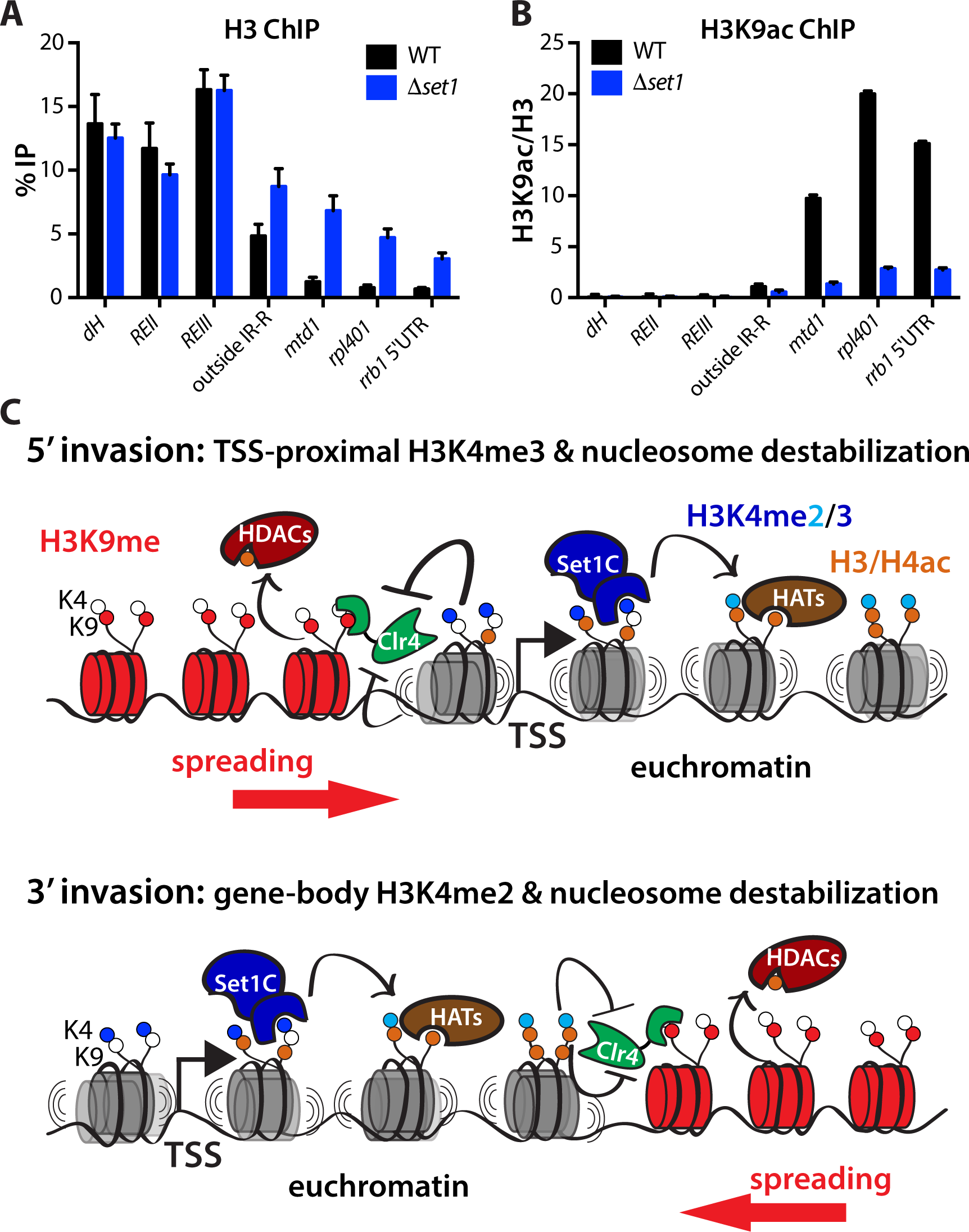
Set1-mediated global destabilization of euchromatic nucleosomes provides a second and orientation-neutral pathway for genic heterochromatin repulsion. **(A)** H3 ChIP in WT (black) and *Δset1* (blue). **(C)** H3K9ac ChIP in WT and *Δset1*. ChIP is normalized to H3 signal to account for differences in nucleosome occupancy. In B. and C. heterochromatin targets include: *dH*, *REII*, and *REIII*. Euchromatic targets include: outside IR-R, *mtd1*, *rpl401*, and *rrb1*. Error bars represent 1SD from four replicates, each representing a single colony deriving from each genotype. **(D)** Model for contribution of Set1/COMPASS to gene-mediated heterochromatin repulsion. In 5′ invasion (TOP), Set1-dependent TSS-proximal H3K4me3 in addition to nucleosome destabilization (reduced occupancy and increased acetylation) counter heterochromatin spreading. In 3′ invasion (BOTTOM) Set1-dependent H3K4me2 in gene bodies and increased nucleosome destabilization counter heterochromatin spreading.

What might lead to this increase in nucleosome occupancy? It is known that histone acetylation is associated with increased nucleosome turnover [24], which, in turn, disrupts heterochromatin stability [24, 33, 60]. Previous studies have identified a role for Set1/COMPASS and H3K4me in promoting global histone acetylation at various residues [38, 61, 62]. To validate this finding in our system, we performed ChIP against H3K9ac, as well as H3 and H4 acetylation broadly, and found that acetylation was similarly reduced in *Δset1* (**Figure 6B, Supplemental Figure 6D,E**). Taken together, the role of Set1/COMPASS in promoting nucleosome destabilization via reduced occupancy and increased acetylation, in addition to catalyzing a kinetic inhibitor of Suv39/Clr4, provide molecular basis for its genome-wide role in containment of heterochromatin spreading at both 3′ and 5′ oriented genes.

## DISCUSSION

Two paradigms have emerged for heterochromatin domain regulation, which when taken together present an intriguing paradox. On one hand is the ability for heterochromatin domains to expand beyond their borders when containment mechanisms are compromised [8, 12, 13, 19, 31, 36, 63]. On the other, is the widespread dispersion of factors, activities, and posttranslational modifications embedded in euchromatin, which are known to antagonize the establishment and maintenance of heterochromatic domains [6, 11, 12, 24, 25, 64]. Why then is heterochromatin spreading able to overcome these negative regulators and expand into euchromatin? Part of the answer may lie in the activities inherently associated with the spreading machinery, including HDACs [64–67], nucleosome remodelers [60, 64], and H3K4 – demethylase complexes [29], which apparently can overpower euchromatin. Yet, how and why heterochromatin spreading is halted at specific euchromatic locations is not understood.

In this work we investigated both the signals within local active euchromatin defines spatial limits to heterochromatin spreading in fission yeast. The key principles that derive from this work are (1) Euchromatic barrier signals depend on Set1/COMPASS activity at active genes. (2) High gene transcript levels are not intrinsically refractory to heterochromatin invasion. (3) Set1*-*dependent repulsion of heterochromatin acts via two pathways, direct catalytic inhibition of Clr4/Suv39 by the Set1-product H3K4me3, and nucleosome mobilization. (4) The ability to repel heterochromatin can be gene orientation specific, likely depending on gene-intrinsic features.

### Regulation of facultative heterochromatin domain size

Regulation of facultative heterochromatin spreading is of critical importance during development as the variable formation of repressed domains directs cell fate decisions [17, 18]. Both H3K27 and H3K9 methylated domains mark genomic regions orthogonal to the intended cell fate and mediate their transcriptional repression. The mechanisms defining the borders of H3K9me-marked domains during development are not well understood. Theoretical work [21] proposed that domain size can be tuned by the ratio of spreading rate and turnover rate (which broadly includes containment), and that above a certain threshold of this ratio, spreading proceeds as if unbounded. While little is known about the potential mechanisms for tuning the rate of spreading, regulation of turnover could result from the directed recruitment of heterochromatin antagonists in response to cellular cues, or the intrinsic placement of modular euchromatic “halt” signals within the path of an expanding domain.

We find Set1/COMPASS enacts a “turnover” or containment signal at gene-rich regions, including facultative heterochromatin in fission yeast (**Figure 2**). These small, euchromatin-embedded, islands of H3K9me form over meiotic genes in response to RNA processing activities [19] and are restricted by the JmjC domain-containing protein Epe1, similarly to constitutive loci. In our sensitized system lacking *epe1*, we identified several heterochromatin islands that expanded in the absence of *set1* (**Figure 2C, Supplemental Figure 2D**). We also document the Set1-sensitivity of the gene-rich region outside the IR-R MAT locus boundary (**Figure 2B**). In contrast to previously identified spreading regulators which function globally, such as Epe1, Leo1, and Mst2 [8–11, 19], the containment function of Set1 is localized to specific euchromatic regions consistent with its role at active genes. In the case of boundary failure, the IR-R proximal region experiences silencing. Silencing of downstream essential genes is likely prevented by the intervening *rpl401* gene, which displays a large peak of H3K4me3 (**Figure 2B)** and strong dependence on *set1* for spreading repulsion (**Figure 3A**). Similarly, the *set1*-sensitive heterochromatin islands are also enriched for peaks of H3K4me3 (**Figure 2C, Supplemental Figure 2D**). However, we also find several islands that do not expand in the absence of Set1. From this we conclude that Set1 largely contributes to the containment of spreading in gene-rich euchromatin although the presence of H3K4me3 is not always predictive. The reasons why some loci are more sensitive than others thus remains to be investigated. Of note, unlike in fission yeast as documented here, in budding yeast, Set1 appears to have a more global heterochromatin-antagonizing role, in concert with H2A.Z [68]. This suggests that locally acting spreading control by Set1 may have co-evolved with H3K9me marked heterochromatin systems, with other factors regulating global control (see above).

### The role of gene orientation in heterochromatin repulsion

The TSS-enriched accumulation of Set1/COMPASS products, especially H3K4me3, naturally raises the question whether the orientation of genes relative to spreading heterochromatin plays a role in containment. Lamina Associated Domains (LADs) are gene-repressive chromatin domains associated with the nuclear periphery that contain both H3K9 and H3K27 methylation (reviewed in [69]). Interestingly, regions immediately flanking LADs are enriched for 5′ oriented genes and concomitant H3K4 methylation [22]. The authors proposed that presence of H3K4me and 5′ oriented genes could help delimit the repressed domain, but no mechanism was explored. Fission yeast lacks a genome-wide bias for proximal-5′-oriented genes at boundaries (**Supplemental Figure 6A**). However, we found that the genic barrier activity of some genes was orientation-sensitive (**Figure 5A**). Such 5′ barrier bias could be explained by the effect H3K4me3 has on Suv39/Clr4 (**Figure 4B**). Contrasting with prior findings [51] we find that Suv39/Clr4 H3K9 methylation catalysis is directly inhibited by Set1 products, most strongly by H3K4me3. This finding represents a rare example of direct regulation of the Suv39/Clr4 SET domain active site, beyond auto-inhibition [70], but is consistent with the effect H3K4me can have on other H3K9 methylases [52–54]. The flipside of an orientation sensitive gene barrier is weaker effectiveness in the other orientation. We describe a mechanism for overcoming 3′ oriented gene barriers via co-transcriptional gene silencing and down-regulation of H3K4me (**Figure 5C-G**), which relieves the potential for catalytic interference. Other genes we examined however were orientation neutral (**Figure 3E, Figure 5B, Supplemental Figure 3A, Supplemental Figure 5G-I**). These genes likely rely on the second mechanism Set1 implements to repel spreading, mobilization of nucleosomes (**Figure 6A, B**, see model below). While we do not fully understand the mechanisms that prevent spreading in a gene-orientation specific manner, we speculate that this may be partially encoded in the gene-intrinsic H3K4me2 and me3 distribution profiles. It is further an attractive hypothesis that 3′ sensitive invasion can provide a tunable spreading mechanism where variable gene silencing is desired, such as in developmental fate decisions.

### Regulation of active and repressed chromatin states by Set1 and COMPASS

How do the mechanisms of heterochromatin regulation we describe for Set1/COMPASS relate its known roles in transcriptional regulation? The recruitment of Set1/COMPASS to chromatin requires H2B monoubiquitination mediated by Rad6 and Bre1 as well as interaction with the Paf1 elongation complex (Paf1C), which engages RNA polymerase and is additionally responsible for activation of Rad6 and Bre1 function on chromatin. Set1/COMPASS also associates with elongating RNA polymerase, giving rise to a characteristic pattern of H3K4 methylation states (see above). Interestingly, previous studies in fission yeast have described a role for Paf1C components Paf1 and Leo1 in antagonizing heterochromatin spreading through promoting increased histone turnover and H4K16 acetylation [9, 10]. Both studies tested, but did not identify, a role for Set1 in their respective systems at loci (IRC1L of the centromere, and IR-L of MAT) where we also do not detect an effect of *Δset1* even in the sensitized *Δepe1* genetic background (**Supplemental Figure 2**).

Several additional data support a model where Set1 and Paf1/Leo1 act in separate pathways to regulate heterochromatin spreading: (1) *set1* appears not to be epistatic to *leo1* in genome-wide genetic interaction study for heterochromatin spreading using an IRC1L reporter [9]. (2) Global H4K16 acetylation levels did not change in response to *Δset1* [38], whereas acetyl marks such as H3K9 and H3K14 were reduced in this background (**Figure 6B** and [38]). (3) In our repelling factor screen, *Δleo1* did not result in the characteristic spreading phenotype seen for Set1/COMPASS complex deletions (data not shown). Taken together these results describe separate mechanisms for spreading regulation by Leo1/Paf1 and Set1/COMPASS.

In contrast to its well-described role at active genes, Set1 has been found to have gene-repressive functions independent of its H3K4me catalytic activity and other members of the COMPASS complex [43]. This non-complex mediated repression functions through interaction with the Clr3 histone deacetylase and recruitment by the Atf1 transcription factor. Consistent with this report, we found that expression of some genes, notably *act1*, increase in the absence of Set1 (**Supplemental Figure 3B**). In contrast to this repressive role for Set1, we repeatedly found that genes in a number of different loci were subjected to heterochromatin spreading in *Δset1* strains (**Figures 1-3, Supplemental Figures 2,3,5**) and that histone acetylation at genes was decreased (**Figure 6B, Supplemental Figure 6D,E**). Additionally, since deletions of five Set1/COMPASS complex components mimicked the *Δset1* spreading containment phenotype (**Supplemental Figure 1**) it is likely that these two functions of Set1 are mediated through different pathways.

### A model for gene-based regulation of heterochromatin spreading by Set1/COMPASS

We propose the following model for how Set1/COMPASS directs heterochromatin containment: Set1-mediated repulsion involves at least two transcription-independent functions, the catalytic inhibition of the H3K9 methylase Suv39/Clr4 (**Figure 4B**) and destabilization of euchromatic nucleosomes (**Figure 6A,B, Supplemental Figure 6B,D,E**). During 5′ invasion, the spreading reaction encounters destabilized nucleosomes as well as the strongly inhibitory promoter-proximal H3K4me3 mark, which repel spreading (**Figure 6C, TOP**). Conversely, during 3′ invasion spreading is similarly challenged with reduced histone occupancy and increased acetylation in addition to the inhibitory H3K4me2 mark decorating the gene body (**Figure 6C, BOTTOM**).

These mechanisms can explain the Set1-dependence of gene-mediated barrier function regardless of orientation. We believe the orientation effect of containment derives from the relative distribution of H3K4me2 and H3K4me3 marks (see above), resulting in more or less polarized Suv39/Clr4 antagonism and nucleosome mobilization. The fact that heterochromatin invasion can overcome these activities when invading from a 3′ direction for some genes by exploiting co-transcriptional gene silencing is an important principle that could be exploited in epigenome engineering approaches.

Not all genes function as effective barriers to spreading, regardless of the presence of H3K4me. Further investigation will be needed to illuminate what properties in addition to Set1 control effectiveness of spreading repulsion. This work lays the ground for future investigation into mechanisms of regulation of heterochromatin domain size in higher order systems which feature complex developmental plans encoded in heterochromatic patterning.

## Acknowledgements

We thank Hiten D Madhani for generous gifts of strains and reagents and use of laboratory equipment. Additionally, we thank Sandra Catania, Michael McManus, and Kieran Mace for helpful discussions on data acquisition, analysis, and interpretation. This work was supported by grants from the National Institutes of Health (DP2GM123484) and the UCSF Program for Breakthrough Biomedical Research (partially funded by the Sandler Foundation) to BA-S, and the ARCS Foundation Scholarship and Hooper Graduate Fellowship to RAG. This work was supported by grants awarded to SB from the German Research Foundation (BR 3511/2-1) and the European Union Network of Excellence EpiGeneSys (HEALTH-2010-257082). SB is Member of the Collaborative Research Center 1064 funded by the German Research Foundation and acknowledges infrastructure support. JEB is a Natural Sciences and Engineering Research Council of Canada postdoctoral fellow. Flow cytometry and FACS data were generated in the UCSF Parnassus Flow Cytometry Core which is supported by the Diabetes Research Center (DRC) grants NIH P30 DK063720 and NIHS10 1S10OD021822-01.

## Materials and Methods

### Strain and plasmid construction

Plasmids used to generate genomic integration constructs were assembled using *in vivo* recombination. *S. pombe* transformants were selected as described [33]. XFP reporters were targeted to specific genomic locations as described [33]. Direct gene knockout constructs were generated using long primer PCR to amplify resistance cassettes with homology to the regions surrounding the open reading frame of the target. Genomic integrations were confirmed by PCR.

### Flow cytometry and FACS sorting

Cells were grown for flow cytometry experiments and as described [33]. Flow cytometry was performed using a Fortessa X20 Dual machine (Becton Dickinson, San Jose, CA) and High Throughput Sampler (HTS) module. Approximately 20,000 to 100,000 cells were collected, dependent on strain growth and volume collected. Fluorescence detection, compensation, and data analysis were as described[33, 34].

For the FACS experiment, cells were grown overnight from OD = 0.05 in YES and in the morning concentrated into a smaller volume (∼3-5x) and filtered with 35–40μm mesh (Corning) to achieve 5-7k events/second on the cytometer and reduce potential for clogs. Cells were first gated for size (forward and side scatter), removal of doublet cells, the presence of the control “red” signal and then sorted into Low and High populations for “orange”. Low “orange” population was defined by signal overlapping a control with no fluors. High “orange” population was defined by signal overlapping the matched background Δ*clr4* control. For each population, 16-18×10^6^ cells were collected for Chromatin Immunoprecipitation and 3×10^6^ cells were collected for RT-qPCR. Cells were processed for downstream analysis immediately following sorting.

### Repelling Factor Screen

An h-reporter strain with “green” and “orange” at the *ura4* locus (*natMX* marked) and “red” at the *leu1* locus (*hygMX* marked) was crossed to a 408-strain subset of the *Bioneer* haploid deletion library (*kanMX* marked). Crosses were performed as described [9, 71] with limited modifications. Briefly, crosses were arrayed onto SPAS plates using a RoToR HDA colony pinning robot (Singer) and mated for 4 days at room temperature. The plates were incubated at 42°C for 4 days following mating to remove haploid and diploid cells, retaining spores. Resultant spores were germinated on YES medium with added Hygromycin B, G418, and nourseothricin for selection of both reporter loci and the appropriate gene deletion. The resultant colonies were passaged into liquid YES and grown overnight for flow cytometry as described above. In the morning, cells were diluted again into YES medium and grown 4-6 hours at 32°C prior to analysis via flow cytometry.

### RNA extraction and quantification

Cells from log phase cultures or FACS sorted cells were pelleted supernatant was decanted, and flash frozen in liquid nitrogen. Pellets were stored at −80°C. RNA extraction was performed as described [33]. cDNA systhesis was performed with either SuperScript RTIII (Invitrogen) and an oligo dT primer (**Supplemental Figure 4C**) or SuperScript RTIV (Invitrogen) and random hexamers (**Figure 5E, Supplemental Figure 3C**) via the manufacturer’s protocol. cDNA samples were quantified by RT-qPCR as described [33]. Values from cDNA targets were normalized to actin for all samples. Samples in **Figure 5E** and **Supplemental Figure 4C** were normalized to the target/actin value for the Δ*clr4* strain of a matched background. For **Supplemental Figure 3C**, given that signal from *act1p* driven “red” increases by ∼50% in Δ*set1* backgrounds, the target/actin values in Δ*set1* samples were multiplied by the mean ratio Δ*set1*/WT of *act1p* driven “red” signal from the 4 WT and mutant pairs in **Supplemental Figure 3B**. This adjusts the normalization for the up-regulation of actin observed in this background.

### Chromatin immunoprecipitation

Chromatin Immunoprecipitation (ChIP) followed by qPCR was performed essentially as described [33] with the following modifications. For **Figure 4D** 16-18×10^6^ cells of both “low” and “high” FACS populations, as well as controls, were collected and processed for ChIP. Prior to lysis, 50×10^6^ cells of independently fixed *S. cerevisiae* W303 strain were added to each population as carrier. ChIP experiments with bulk populations of log phase cells were performed as described [33] without the addition of W303 carrier. In **Supplemental Figure 6B**, Hht2-HA cells were grown at 25°C, 225rpm in YES+Hygromycin B from OD=0.05. After cells reached OD=0.2, G2 stall was induced by shifting the temperature to 37°C for 3 hours prior to fixation. Following lysis, sonication was performed using the Diagenode BioRuptor Pico machine for 20-28 rounds of 30s ON/30s rest. Cleared chromatin was split into equal volumes per IP after a small fraction (5-10%) was set aside as Input/WCE. 1μL of the following antibodies were added per ChIP sample: H3K9me2 (Abcam ab1220), H3K4me3 (Active Motif 39159), H3K9ac (Active Motif 39137), H3(pan)ac (Active Motif 39064), H4(pan)ac (Active Motif 39140), HA (Abcam ab9110). 1.4μg of H3 antibody (Active Motif 39064) was added per ChIP sample. DNA was quantified by RT-qPCR and %IP (ChIP DNA / Input DNA) was calculated as described [33].

### ChIP-Seq Sample and Library Preparation

Sample preparation and ChIP prior to sequencing was performed essentially as described [33] with the following modifications. 50mL of cells were grown to OD=0.6-0.8 overnight from OD=0.025. Biological duplicate samples were generated for WT, biological triplicate samples were generated for Δ*epe1*, and four biological samples were generated for Δ*epe1*Δ*set1* genotypes. Base on OD measurements, 300×10^6^ cells per sample were fixed and processed for ChIP. Shearing was performed with 20 cycles of 30s ON/30s rest. Samples were not pre-cleared. Sonication efficiency was determined for each sample and only samples where DNAs averaged 200-300bp were used. Chromatin was split into two samples after 8% was set aside as input. 3μL of H3K9me2 (abcam1220) or H3K4me3 (Active Motif 39159) antibodies were added per tube and incubated overnight at 4°C with rotation. (Only H3K9me2 ChIP was performed for Δ*set1* strains. The absence of H3K4me3 was validated by ChIP qPCR in **Supplemental Figure 2E**). Immune complexes were collected with 30μL twice-washed Protein A Dynabeads (Invitrogen) for 3 hours at 4°C. Beads were washed as above with the exception that the Wash Buffer step was performed twice. Following incubation at 70°C for 20 minutes, DNA was eluted in 100μl of TE + 1%SDS and the beads were washed and eluted a second time with 100μl of TE + 1%SDS + 5 μl of 20mg/mL Proteinase K (Roche). Following overnight incubation at 65°C, ChIP and Input samples were purified using Machery Nagel PCR clean up kit. Library preparation for sequencing was performed as described [72, 73]. Samples were sequenced on a HiSeq 4000 platform (Illumina) with a Single End 50 run.

### ChIP-seq data analysis

Sliding window quality filtering and adapter trimming were carried out using Trimmomatic 0.38 [74] before the reads were aligned to the *S. pombe* genome [75] with Bowtie2 2.3.4.2 [76] using standard end-to-end sensitive alignment. Indexed bam files were generated using SAMtools 1.9 [77] “view”, “sort”, and “index” functions. Combined Input files and WT H3K9me2 ChIP files were generated with SAMtools “merge” function for use in normalization. Input or WT normalized signal tracks were generated using the MACS2 version 2.1.1.20160309 [78, 79] callpeak function to generate reads per million normalized bedGraph files with the following flags: -g 1.26e7 --nomodel --extsize 200 --keep-dup auto -B --SPMR -q 0.01. The resulting pileup was normalized with the bdgcmp function via the fold enrichment method (m –FE). The resulting normalized signal track files were trimmed back to the length of the genome and converted to bigwig format using UCSCtools bedClip and bedGraphToBigWig functions. BigWig files were imported into R 3.5.1 with rtracklayer 1.40.6 [80]. The genome was divided into 25bp bins and the average enrichment value per bin was calculated using the tileGenome and binnedAverage functions of GenomicRanges 1.32.7 [81]. Gene annotations were imported from PomBase [82] and converted to genomic coordinates with the makeTxDbFromGFF function from GenomicFeatures 1.32.3 [81]. Finally mean and confidence interval per each genotype were generated during signal track plotting using the DataTrack command from Gviz 1.24.0 [83]. Peaks were called with epic2 0.0.14 [84] with the following flags: --effective-genome-fraction 0.999968 -bin 200 -g 3 -fs 200 –fdr 0.05. Regions of known heterochromatin formation were imported from a previously curated list [72]. Regions were extended by 10kb on each side to account for differences in coordinates that may exist for different genome assemblies, as well as variable spreading. Peaks and known regions were plotted using Gviz [83].

### Clr4 Chromodomain and Clr4 SET domain Purification

The chromodomain of Clr4 (residues 6-64, Clr4-CD) and SET domain (residues 192-490, Clr4-SET) were each cloned into MacroLab vector 14C containing N-terminal 6xHis and Maltose Binding Protein (MBP) tags. Proteins were expressed as described [48] except that for Clr4-SET, LB was substituted for 2XYT medium supplemented with 10μM ZnSO_4_. Lysis and Talon affinity resin purification (Takara Bio) and size exclusion chromatography was essentially as described [48]. Lysis Buffer was 100mM HEPES pH 7.5, 300mM NaCl, 10% glycerol, 7.5mM imidazole, 0.5% Triton-X100, 1mM β-mercaptoethanol, and protease inhibitors. For Clr4-SET, Triton was substituted for 0.01% Igepal NP-40. After final size exclusion chromatography, Clr4-CD was eluted into FP storage buffer (20mM HEPES pH 7.5, 100mM KCl, 10% glycerol, and 5mM β-mercaptoethanol). Clr4-SET was eluted into Clr4 Storage Buffer (100 mM Tris pH 8.5, 100 mM KCl, 10% glycerol, 1 mM MgCl_2_, 20µM ZnSO_4_, and 10 mM β-mercaptoethanol). All proteins were flash frozen and stored at −80°C. Protein concentration was determined by Sypro Ruby (Biorad) gel staining against a BSA standard curve and verified by UV absorption at 280 nm using the theoretical extinction coefficient (ExPasy ProtParam) 88810cm^-1^M^-1^ and 98210cm^-1^M^-1^ for Clr4-CD and Clr4-SET, respectively.

### Fluorescence Polarization Assay

Fluorescence polarization assay for binding of Clr4-CD to H3 tail peptides was performed as described [85]. 10nM of H3 tail peptide with K4me0K9me0 (unmodified), K4me0K9me3, or K4me3K9me3 modifications (GenScript) was use as probe. Reactions were performed in FP buffer (20mM HEPES pH 7.5, 100mM KCl, 10% glycerol, and 0.01% NP-40 substitute), and incubated for 20 minutes at RT prior to measurement. Fluorescence polarization measurements and data analysis including fitting of curves were performed as described [85].

### Histone Methyltransferase Assay

Multiple turnover kinetic assay was performed as described [48] with the following modifications. Reactions contained 100 µM cold SAM (disulfate tosylate, Abcam) and 10-15µM ^3H^SAM tracer (55–75 Ci/mmol, PerkinElmer) and were incubated with 1µM Suv39/Clr4-SET and varying amounts of biotinylated H3(1–20) peptide with K4me0 (unmodified), K4me2, or K4me3 (GenScript). Reactions were performed at 30°C in Clr4 Reaction Buffer (100-120 mM Tris pH 8.5, 100 mM KCl, 10% glycerol, 1 mM MgCl2, 20 µM ZnSO4, and 10 mM β -mercaptoethanol).

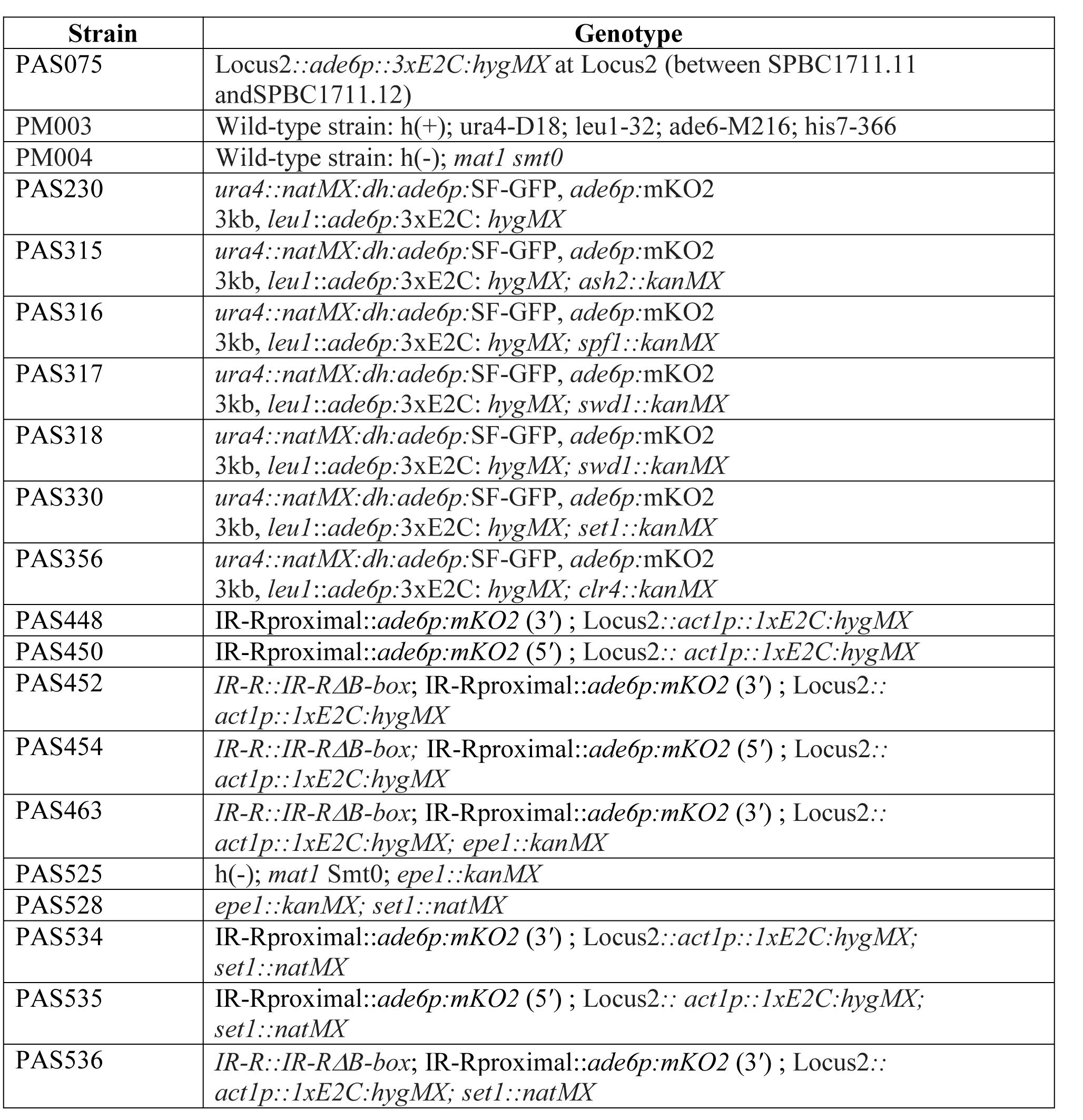
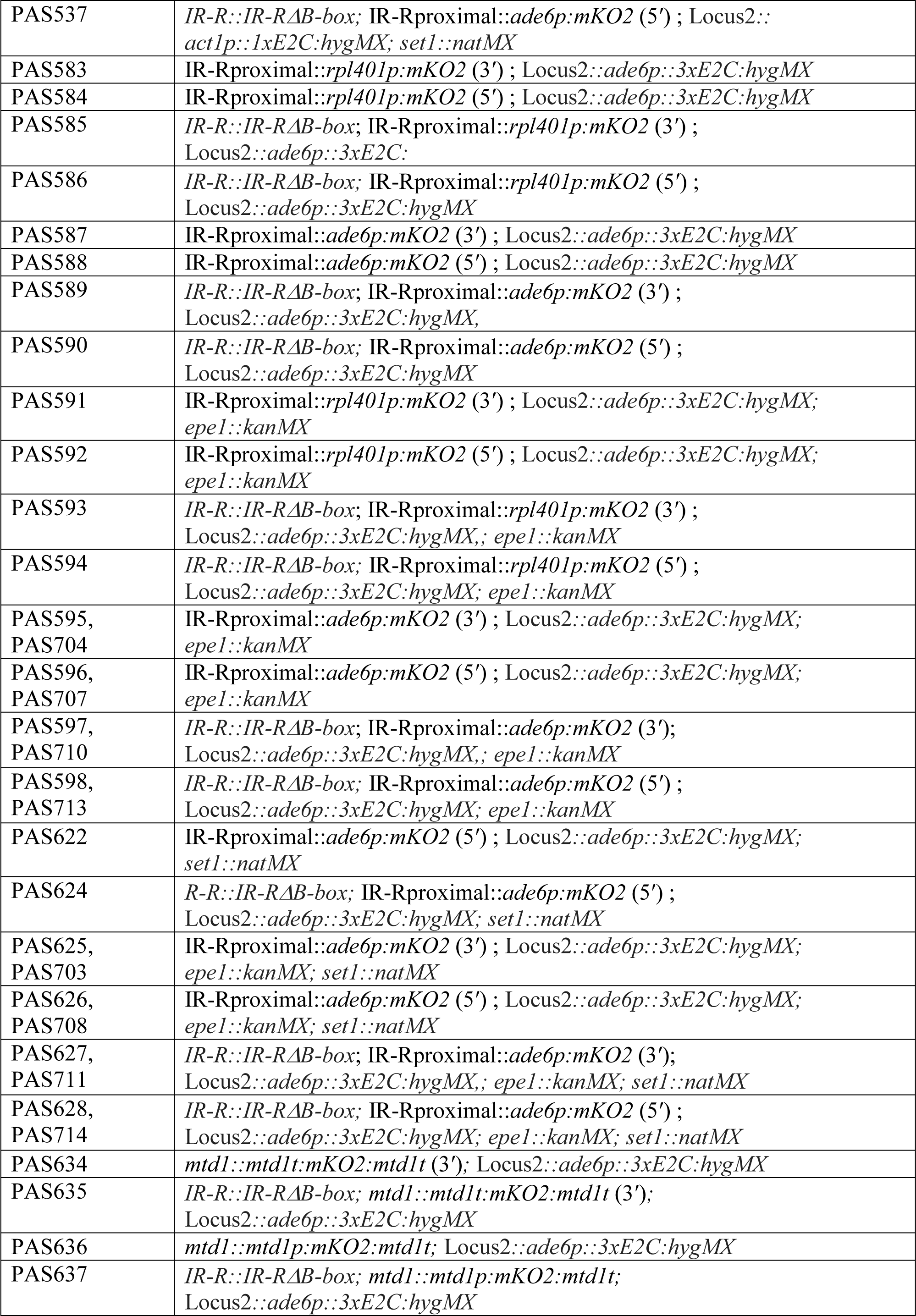
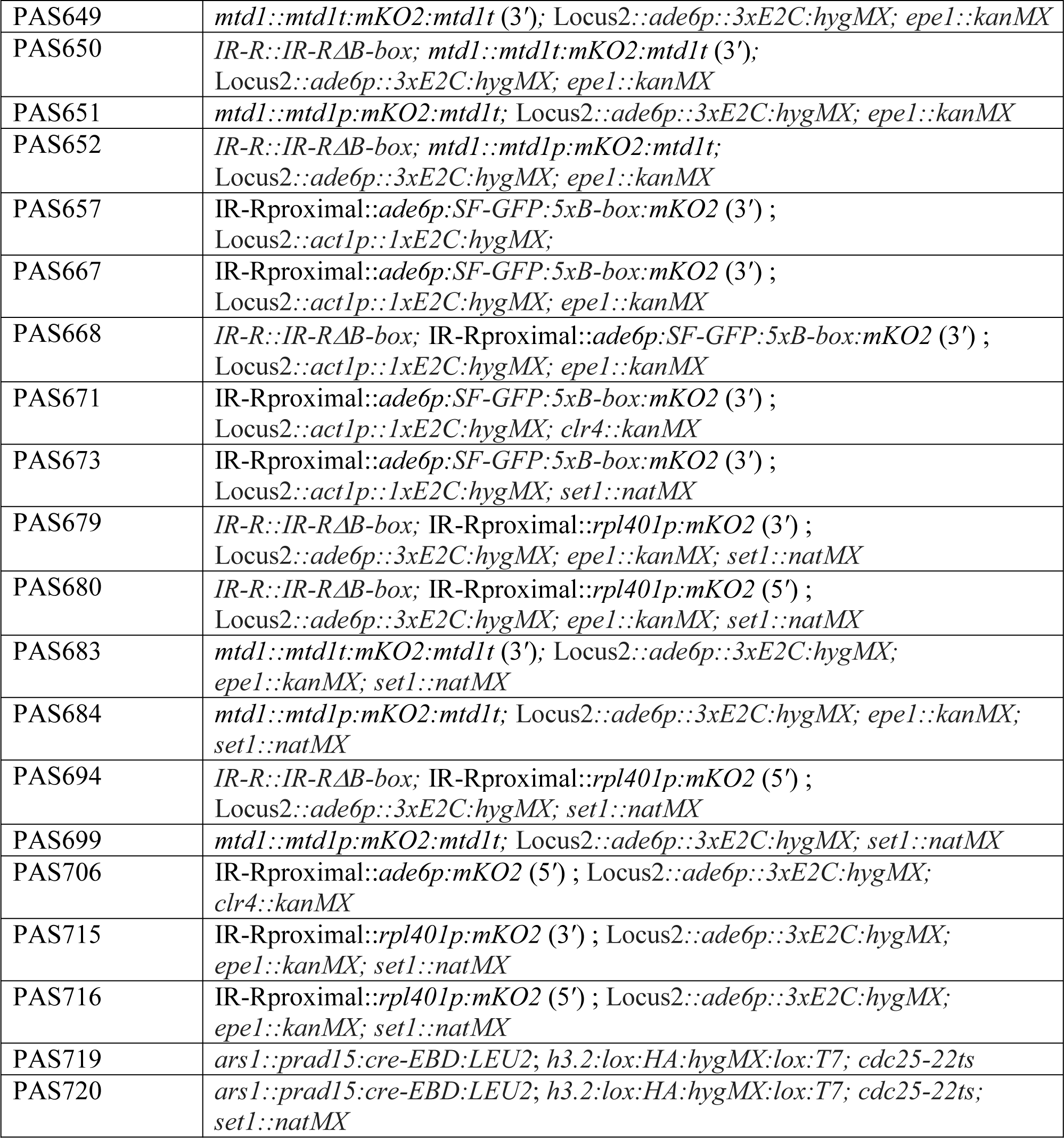

---

**Supplemental Figure 1:**
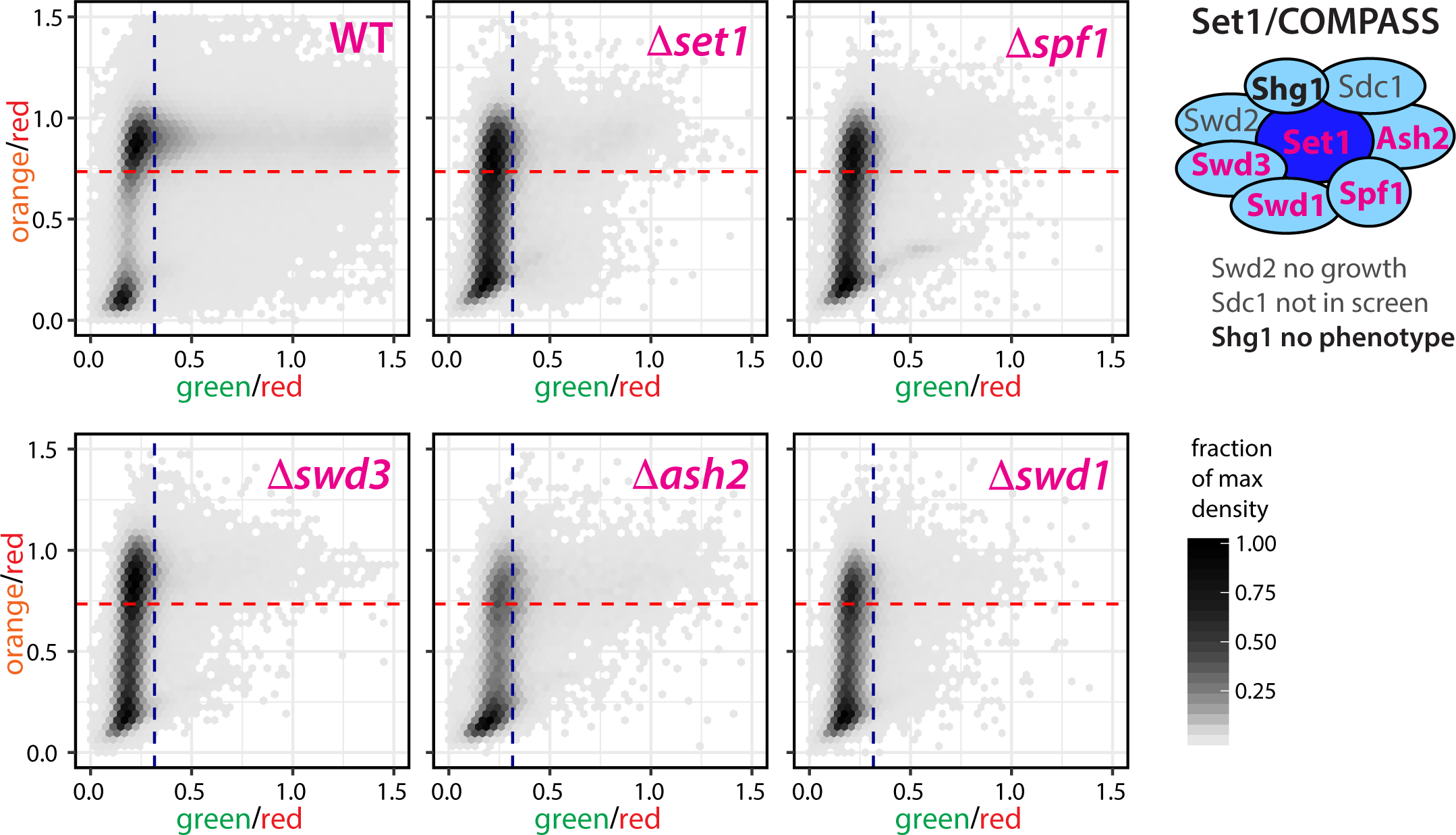
Set1/COMPASS regulates gene-mediated heterochromatin barriers. Two dimensional density hexbin plots of red-normalized “green” and “orange” signal from WT parental strain and deletions of five Set1/COMPASS complex members. Cells with values below the nucleation cutoff (blue line) and spreading cutoff (red line) are considered repressed.

**Supplemental Figure 2:**
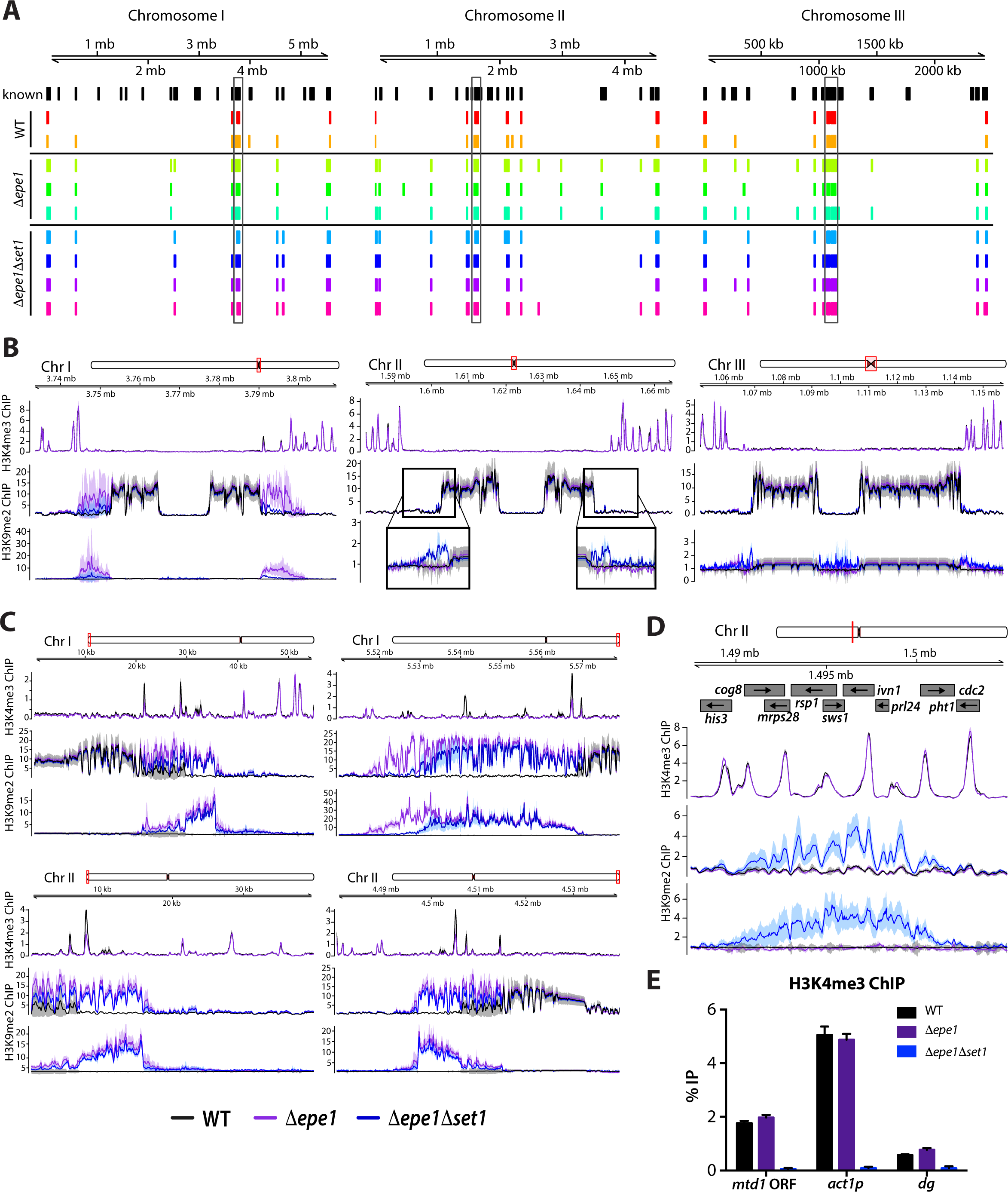
The effect of Epe1 and Set1 at constitutive and facultative heterochromatin loci. **(A)** The location of H3K9me2 peaks in each strain on each chromosome. Previously identified H3K9me peaks [11, 19, 72, 86] were included as reference (black). Regions were extended by 10kb on each side to account for differences in coordinates that may exist for different genome assemblies, as well as variable spreading. Centromeric regions are boxed in grey. **(B)** ChIP-Seq signal tracks plotted in Figure 2 for centromere proximal regions. H3K9me2 ChIP normalized to WT for centromere II is cropped and expanded for a closer view. **(C)** ChIP-Seq signal tracks for telomere proximal regions of chromosomes I and II. **(D)** ChIP-Seq signal tracks for novel heterochromatin region specific to the *Δepe1Δset1* genotype. **(E)** ChIP-qPCR data to validate the absence of H3K4 methylation in *Δset1;* Error bars represent 1SD from three technical replicate ChIPs.

**Supplemental Figure 3:**
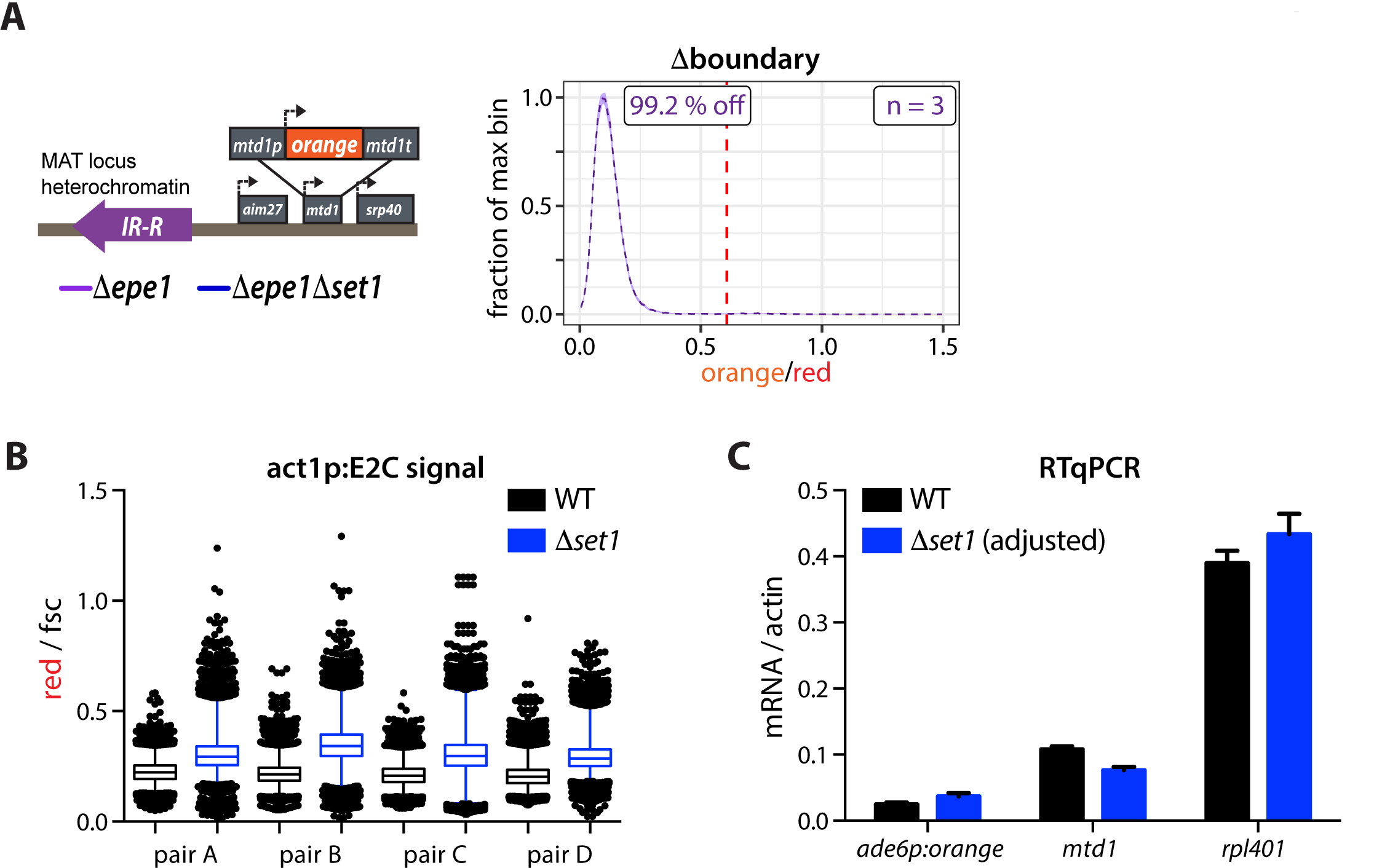
Set1 represses transcription of *act1*, however RNA levels of boundary associated genes are not broadly affected. (A) Overview of the *mtd1p*-HSS. (B) Histogram plots as in Figure 1 of normalized orange signal from *set1*+ (purple) *mtd1p*-HSS Δboundary isolates. (C) Box and whisker plots of fsc normalized “red” signal plotted as in Figure 3F for four independent pairs of *set1*+ (black) and *Δset1* (blue). (C) RT-qPCR signal for the *ade6p*-driven “orange” transcript, as well as native *mtd1* and *rpl401* transcripts. Values are normalized to signal from *act1*/actin. In *Δset1* the *act1* signal was adjusted for the mean ratio of *act1* in *Δset1* to *set1*+ as seen in B. Error bars indicate standard deviation of 3 technical replicate RNA isolations.

**Supplemental Figure 4:**
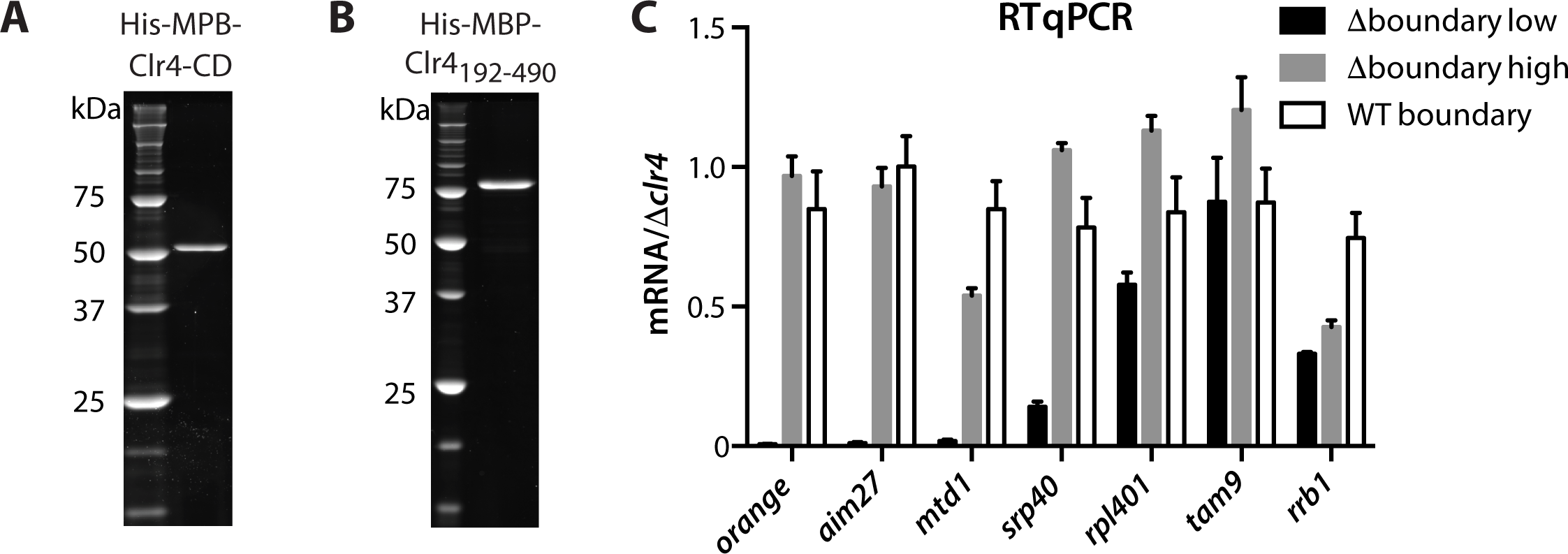
Gene-protective activity of Set1/COMPASS is rooted in catalytic inhibition of Suv39/Clr4 by H3K4me2/3 (A) SDS-PAGE gel with His-MBP-Clr4-CD used in Figure 4A. (B) SDS-PAGE gel with His-MBP-Clr4-192-490 used in Figure 4B. (C) RT-qPCR data from “low” and “high” sorted populations and one WT isolate from Figure 5A,B. Signal is normalized to values from corresponding *Δclr4* strain. Error bars represent 1SD of three technical replicates.

**Supplemental Figure 5:**
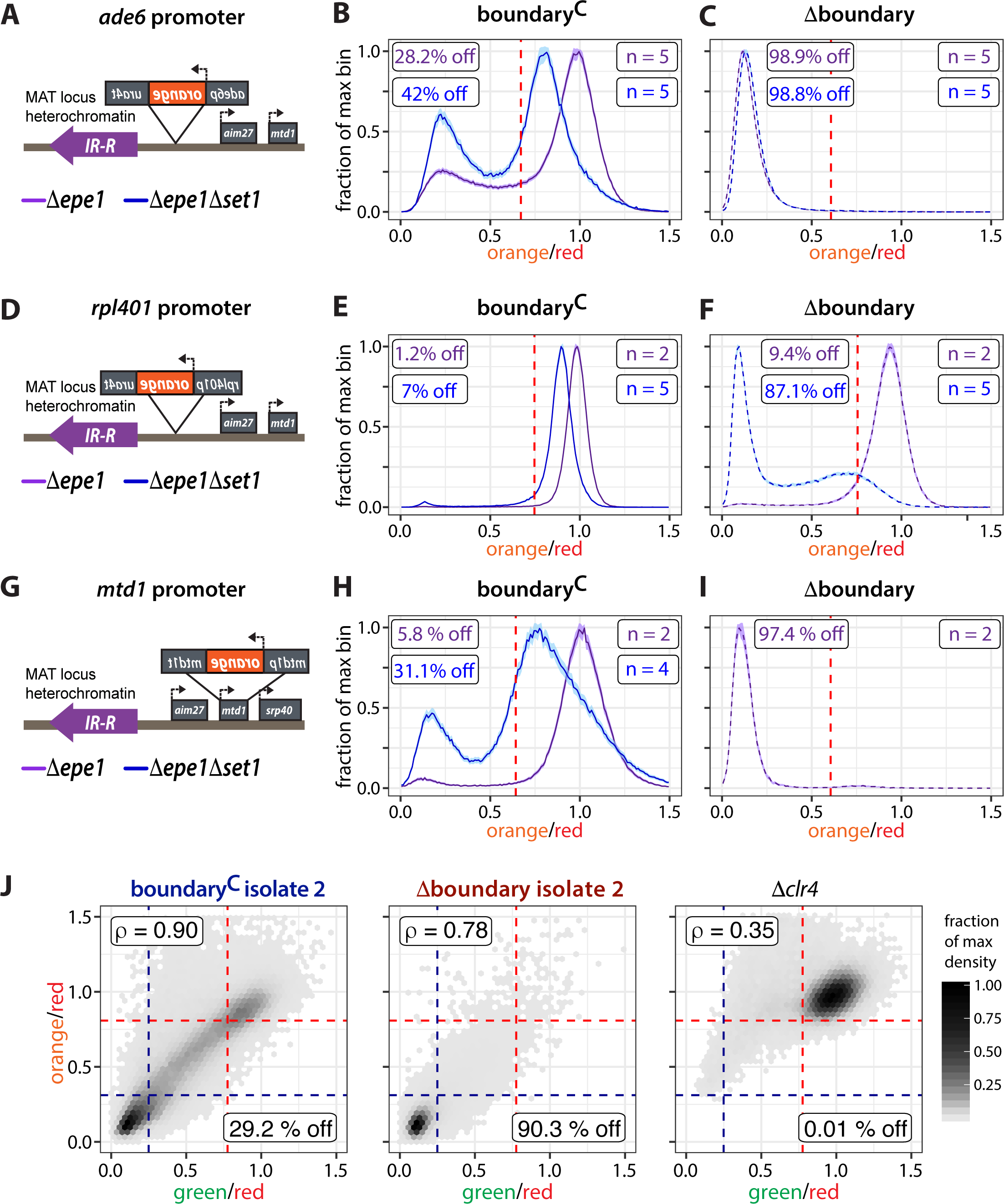
Response of 3*′* oriented genes to heterochromatin spreading. **(A)** Locus cartoon for 3′ *ade6p*-driven “orange”. **(B)** Histogram of normalized *ade6p-* “orange” signal in boundary^C^ in *set1+* (purple) and *Δset1* (blue) background as in Figure 1. **(C)** Histogram of normalized *ade6p-* “orange” signal in Δboundary strains. **(D)** Locus cartoon for 3′ *rpl401p*-driven “orange”. **(E)** Histogram of normalized *rpl401p-* “orange” signal in boundary^C^ strains. **(F)** Histogram of normalized *rpl401p-* “orange” signal in Δboundary strains. **(G)** Locus cartoon for 3′ *mtd1p*-driven “orange”. **(H)** Histogram of normalized *mtd1p-* “orange” signal in boundary^C^ strains **(I)** Histogram of normalized *mtd1p-* “orange” signal in Δboundary in *set1+* (purple) strains. **(J)** boundary^C^ and Δboundary isolate 2 and well as *Δclr4* strains from Figure 5D-G plotted as in Figure 5D.

**Supplemental Figure 6:**
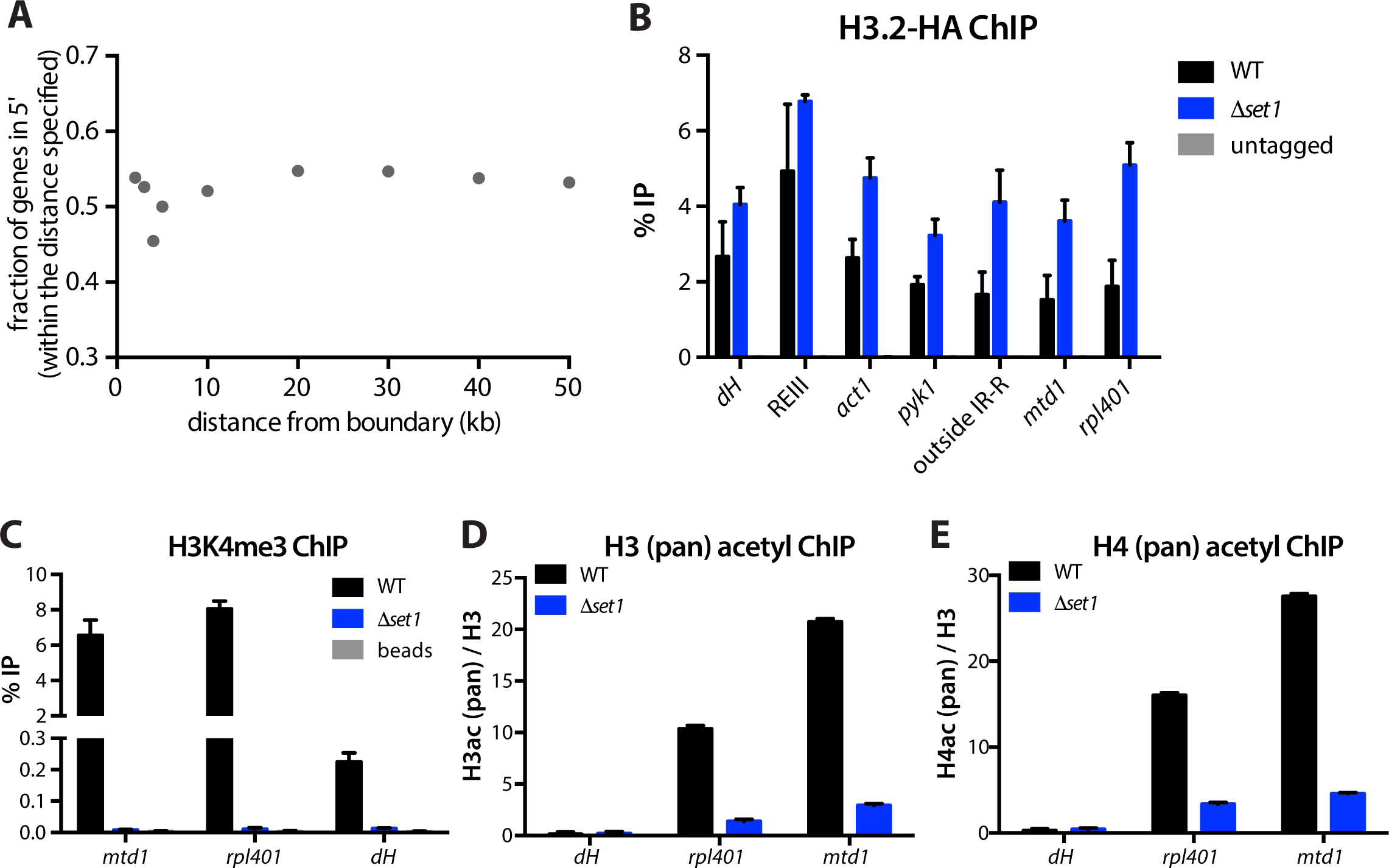
Set1-mediated global destabilization of euchromatic nucleosomes provides a basis for orientation-neutral distribution of boundary-proximal genes in fission yeast. (A) Analysis of gene orientation adjacent to canonical heterochromatin boundaries (centromeres and MAT locus). The fraction of genes within the distance specific (x-axis) in the 5′ orientation with respect to boundaries is plotted. (B) HA ChIP signal from WT (black) and *Δset1* (blue) strains. In these cells, the *hht2* was tagged with an HA epitope and they expressed the *cdc25-22ts* allele. Cells were stalled in G2 phase by shifting the temperature to 37°C for three hours prior to fixation. Error bars represent 1SD from three biological replicates. (C) H3K4me3 ChIP for the cells represented in Figure 6A,B validating the absence of *set1.* Error bars represent 1SD from four biological replicates. (D) H3 (pan) Acetyl ChIP plotted as in Figure 6C. Error bars represent 1SD from four biological replicates. (E) H4 (pan) Acetyl ChIP plotted as in Figure 6D. Error bars represent 1SD from four biological replicates.

## References

1. Rea, S., et al., Regulation of chromatin structure by site-specific histone H3 methyltransferases. Nature, 2000. 406: p. 593.

2. Hall, I.M., et al., Establishment and maintenance of a heterochromatin domain. Science, 2002. 297(5590): p. 2232-7.

3. Jia, S., K.-i. Noma, and S.I.S. Grewal, RNAi-Independent Heterochromatin Nucleation by the Stress-Activated ATF/CREB Family Proteins. Science, 2004. 304(5679): p. 1971.

4. Reyes-Turcu, F.E., et al., Defects in RNA quality control factors reveal RNAi-independent nucleation of heterochromatin. Nature Structural & Molecular Biology, 2011. 18: p. 1132.

5. Braun, S., et al., The Cul4-Ddb1(Cdt2) Ubiquitin Ligase Inhibits Invasion of a Boundary-Associated Antisilencing Factor into Heterochromatin. Cell, 2011. 144(1): p. 41-54.

6. Lan, F., et al., S. pombe LSD1 Homologs Regulate Heterochromatin Propagation and Euchromatic Gene Transcription. Molecular Cell, 2007. 26(1): p. 89-101.

7. Ayoub, N., et al., A novel jmjC domain protein modulates heterochromatization in fission yeast. Mol Cell Biol, 2003. 23(12): p. 4356-70.

8. Trewick, S.C., et al., The JmjC domain protein Epe1 prevents unregulated assembly and disassembly of heterochromatin. The EMBO Journal, 2007. 26(22): p. 4670-4682.

9. Verrier, L., et al., Global regulation of heterochromatin spreading by Leo1. Open Biol, 2015. 5(5).

10. Sadeghi, L., et al., The Paf1 complex factors Leo1 and Paf1 promote local histone turnover to modulate chromatin states in fission yeast. EMBO reports, 2015. 16(12): p. 1673-1687.

11. Wang, J., B.D. Reddy, and S. Jia, Rapid epigenetic adaptation to uncontrolled heterochromatin spreading. eLife, 2015. 4: p. e06179.

12. Wang, J., et al., Epe1 recruits BET family bromodomain protein Bdf2 to establish heterochromatin boundaries. Genes & Development, 2013. 27(17): p. 1886-1902.

13. Noma, K., et al., A role for TFIIIC transcription factor complex in genome organization. Cell, 2006. 125(5): p. 859-72.

14. Bell, A.C., A.G. West, and G. Felsenfeld, The protein CTCF is required for the enhancer blocking activity of vertebrate insulators. Cell, 1999. 98(3): p. 387-96.

15. Kurukuti, S., et al., CTCF binding at the *H19* imprinting control region mediates maternally inherited higher-order chromatin conformation to restrict enhancer access to *Igf2*. Proceedings of the National Academy of Sciences, 2006. 103(28): p. 10684.

16. Ceol, C.J., et al., The histone methyltransferase SETDB1 is recurrently amplified in melanoma and accelerates its onset. Nature, 2011. 471(7339): p. 513-7.

17. Wen, B., et al., Large histone H3 lysine 9 dimethylated chromatin blocks distinguish differentiated from embryonic stem cells. Nat Genet, 2009. 41(2): p. 246-50.

18. Zhu, J., et al., Genome-wide Chromatin State Transitions Associated with Developmental and Environmental Cues. Cell, 2013. 152(3): p. 642-654.

19. Zofall, M., et al., RNA elimination machinery targeting meiotic mRNAs promotes facultative heterochromatin formation. Science (New York, N.Y.), 2012. 335(6064): p. 96-100.

20. McDonald, O.G., et al., Genome-scale epigenetic reprogramming during epithelial-to-mesenchymal transition. Nat Struct Mol Biol, 2011. 18(8): p. 867-74.

21. Hathaway, N.A., et al., Dynamics and memory of heterochromatin in living cells. Cell, 2012. 149(7): p. 1447-60.

22. Guelen, L., et al., Domain organization of human chromosomes revealed by mapping of nuclear lamina interactions. Nature, 2008. 453: p. 948.

23. Scott, K.C., S.L. Merrett, and H.F. Willard, A heterochromatin barrier partitions the fission yeast centromere into discrete chromatin domains. Curr Biol, 2006. 16(2): p. 119-29.

24. Aygün, O., S. Mehta, and S.I.S. Grewal, HDAC-mediated suppression of histone turnover promotes epigenetic stability of heterochromatin. Nature structural & molecular biology, 2013. 20(5): p. 547-554.

25. Garcia, J.F., et al., Combinatorial, site-specific requirement for heterochromatic silencing factors in the elimination of nucleosome-free regions. Genes & development, 2010. 24(16): p. 1758-1771.

26. Noma, K., C.D. Allis, and S.I.S. Grewal, Transitions in distinct histone H3 methylation patterns at the heterochromatin domain boundaries. Science, 2001. 293(5532): p. 1150-1155.

27. Cam, H.P., et al., Comprehensive analysis of heterochromatin- and RNAi-mediated epigenetic control of the fission yeast genome. Nature Genetics, 2005. 37: p. 809.

28. Litt, M.D., et al., Correlation Between Histone Lysine Methylation and Developmental Changes at the Chicken β-Globin Locus. Science, 2001. 293(5539): p. 2453.

29. Li, F., et al., Lid2, a JmjC domain-Containing Histone Demethylase, is Required for Coordinating H3K4 and H3K9 Methylation of Heterochromatin and Euchromatin. Cell, 2008. 135(2): p. 272-283.

30. Noma, K.-i., et al., A Role for TFIIIC Transcription Factor Complex in Genome Organization. Cell, 2006. 125(5): p. 859-872.

31. Zofall, M. and S.I. Grewal, Swi6/HP1 recruits a JmjC domain protein to facilitate transcription of heterochromatic repeats. Mol Cell, 2006. 22(5): p. 681-92.

32. Shilatifard, A., The COMPASS family of histone H3K4 methylases: mechanisms of regulation in development and disease pathogenesis. Annual review of biochemistry, 2012. 81: p. 65-95.

33. Greenstein, R.A., et al., Noncoding RNA-nucleated heterochromatin spreading is intrinsically labile and requires accessory elements for epigenetic stability. eLife, 2018. 7: p. e32948.

34. Al-Sady, B., et al., Sensitive and Quantitative Three-Color Protein Imaging in Fission Yeast Using Spectrally Diverse, Recoded Fluorescent Proteins with Experimentally-Characterized In Vivo Maturation Kinetics. PLoS ONE, 2016. 11(8): p. e0159292.

35. Ayoub, N., et al., A fission yeast repression element cooperates with centromere-like sequences and defines a mat silent domain boundary. Genetics, 2000. 156(3): p. 983-94.

36. Garcia, J.F., B. Al-Sady, and H.D. Madhani, Intrinsic Toxicity of Unchecked Heterochromatin Spread Is Suppressed by Redundant Chromatin Boundary Functions in *Schizosacchromyces pombe*. G3: Genes|Genomes|Genetics, 2015. 5(7): p. 1453.

37. Marina, D.B., et al., A conserved ncRNA-binding protein recruits silencing factors to heterochromatin through an RNAi-independent mechanism. Genes & Development, 2013. 27(17): p. 1851-1856.

38. Noma, K. and S.I. Grewal, Histone H3 lysine 4 methylation is mediated by Set1 and promotes maintenance of active chromatin states in fission yeast. Proc Natl Acad Sci U S A, 2002. 99 Suppl 4: p. 16438-45.

39. Miller, T., et al., COMPASS: a complex of proteins associated with a trithorax-related SET domain protein. Proceedings of the National Academy of Sciences of the United States of America, 2001. 98(23): p. 12902-12907.

40. Roguev, A., et al., High conservation of the Set1/Rad6 axis of histone 3 lysine 4 methylation in budding and fission yeasts. J Biol Chem, 2003. 278(10): p. 8487-93.

41. Santos-Rosa, H., et al., Active genes are tri-methylated at K4 of histone H3. Nature, 2002. 419: p. 407.

42. Marguerat, S., et al., Quantitative analysis of fission yeast transcriptomes and proteomes in proliferating and quiescent cells. Cell, 2012. 151(3): p. 671-83.

43. Lorenz, D.R., et al., Heterochromatin assembly and transcriptome repression by Set1 in coordination with a class II histone deacetylase. eLife, 2014. 3: p. e04506.

44. Mikheyeva, I.V., et al., Multifaceted Genome Control by Set1 Dependent and Independent of H3K4 Methylation and the Set1C/COMPASS Complex. PLoS Genetics, 2014. 10(10): p. e1004740.

45. Buratowski, S. and T. Kim, The role of cotranscriptional histone methylations. Cold Spring Harbor symposia on quantitative biology, 2010. 75: p. 95-102.

46. D’Urso, A., et al., Set1/COMPASS and Mediator are repurposed to promote epigenetic transcriptional memory. Elife, 2016. 5.

47. Muller, M.M., et al., A two-state activation mechanism controls the histone methyltransferase Suv39h1. Nat Chem Biol, 2016. 12(3): p. 188-93.

48. Al-Sady, B., H.D. Madhani, and G.J. Narlikar, Division of labor between the chromodomains of HP1 and Suv39 methylase enables coordination of heterochromatin spread. Mol Cell, 2013. 51(1): p. 80-91.

49. Margueron, R., et al., Role of the polycomb protein EED in the propagation of repressive histone marks. Nature, 2009. 461(7265): p. 762-767.

50. Xhemalce, B. and T. Kouzarides, A chromodomain switch mediated by histone H3 Lys 4 acetylation regulates heterochromatin assembly. Genes & Development, 2010. 24(7): p. 647-652.

51. Nakayama, J., et al., Role of histone H3 lysine 9 methylation in epigenetic control of heterochromatin assembly. Science, 2001. 292(5514): p. 110-3.

52. Wang, H., et al., Purification and Functional Characterization of a Histone H3-Lysine 4-Specific Methyltransferase. Molecular Cell, 2001. 8(6): p. 1207-1217.

53. Nishioka, K., et al., Set9, a novel histone H3 methyltransferase that facilitates transcription by precluding histone tail modifications required for heterochromatin formation. Genes & development, 2002. 16(4): p. 479-489.

54. Binda, O., et al., Trimethylation of histone H3 lysine 4 impairs methylation of histone H3 lysine 9: Regulation of lysine methyltransferases by physical interaction with their substrates. Epigenetics, 2010. 5(8): p. 767-775.

55. Collazo, E., et al., A coupled fluorescent assay for histone methyltransferases. Analytical Biochemistry, 2005. 342(1): p. 86-92.

56. Dirk, L.M.A., et al., Kinetic Manifestation of Processivity during Multiple Methylations Catalyzed by SET Domain Protein Methyltransferases. Biochemistry, 2007. 46(12): p. 3905-3915.

57. Barski, A., et al., High-resolution profiling of histone methylations in the human genome. Cell, 2007. 129(4): p. 823-37.

58. Pokholok, D.K., et al., Genome-wide Map of Nucleosome Acetylation and Methylation in Yeast. Cell, 2005. 122(4): p. 517-527.

59. Liu, C.L., et al., Single-nucleosome mapping of histone modifications in S. cerevisiae. PLoS Biol, 2005. 3(10): p. e328.

60. Taneja, N., et al., SNF2 Family Protein Fft3 Suppresses Nucleosome Turnover to Promote Epigenetic Inheritance and Proper Replication. Mol Cell, 2017. 66(1): p. 50-62.e6.

61. Ginsburg, D.S., et al., NuA4 links methylation of histone H3 lysines 4 and 36 to acetylation of histones H4 and H3. The Journal of biological chemistry, 2014. 289(47): p. 32656-32670.

62. Taverna, S.D., et al., Yng1 PHD finger binding to H3 trimethylated at K4 promotes NuA3 HAT activity at K14 of H3 and transcription at a subset of targeted ORFs. Mol Cell, 2006. 24(5): p. 785-796.

63. Wang, J., et al., Chromosome boundary elements and regulation of heterochromatin spreading. Cellular and molecular life sciences: CMLS, 2014. 71(24): p. 4841-4852.

64. Sugiyama, T., et al., SHREC, an effector complex for heterochromatic transcriptional silencing. Cell, 2007. 128(3): p. 491-504.

65. Yamada, T., et al., The nucleation and maintenance of heterochromatin by a histone deacetylase in fission yeast. Mol Cell, 2005. 20(2): p. 173-85.

66. Shankaranarayana, G.D., et al., Sir2 regulates histone H3 lysine 9 methylation and heterochromatin assembly in fission yeast. Curr Biol, 2003. 13(14): p. 1240-6.

67. Grewal, S.I., M.J. Bonaduce, and A.J. Klar, Histone deacetylase homologs regulate epigenetic inheritance of transcriptional silencing and chromosome segregation in fission yeast. Genetics, 1998. 150(2): p. 563-76.

68. Venkatasubrahmanyam, S., et al., Genome-wide, as opposed to local, antisilencing is mediated redundantly by the euchromatic factors Set1 and H2A.Z. Proc Natl Acad Sci U S A, 2007. 104(42): p. 16609-14.

69. van Steensel, B. and A.S. Belmont, Lamina-Associated Domains: Links with Chromosome Architecture, Heterochromatin, and Gene Repression. Cell, 2017. 169(5): p. 780-791.

70. Iglesias, N., et al., Automethylation-induced conformational switch in Clr4 (Suv39h) maintains epigenetic stability. Nature, 2018. 560(7719): p. 504-508.

71. Barrales, R.R., et al., Control of heterochromatin localization and silencing by the nuclear membrane protein Lem2. Genes Dev, 2016. 30(2): p. 133-48.

72. Parsa, J.-Y., et al., Polymerase pausing induced by sequence-specific RNA-binding protein drives heterochromatin assembly. Genes & Development, 2018. 32(13-14): p. 953-964.

73. Inada, M., et al., Phospho-site mutants of the RNA Polymerase II C-terminal domain alter subtelomeric gene expression and chromatin modification state in fission yeast. Nucleic Acids Res, 2016. 44(19): p. 9180-9189.

74. Bolger, A.M., M. Lohse, and B. Usadel, Trimmomatic: a flexible trimmer for Illumina sequence data. Bioinformatics (Oxford, England), 2014. 30(15): p. 2114-2120.

75. Wood, V., et al., The genome sequence of Schizosaccharomyces pombe. Nature, 2002. 415(6874): p. 871-80.

76. Langmead, B. and S.L. Salzberg, Fast gapped-read alignment with Bowtie 2. Nature methods, 2012. 9(4): p. 357-359.

77. Li, H., et al., The Sequence Alignment/Map format and SAMtools. Bioinformatics, 2009. 25(16): p. 2078-9.

78. Feng, J., et al., Identifying ChIP-seq enrichment using MACS. Nat Protoc, 2012. 7(9): p. 1728-40.

79. Zhang, Y., et al., Model-based analysis of ChIP-Seq (MACS). Genome Biol, 2008. 9(9): p. R137.

80. Lawrence, M., R. Gentleman, and V. Carey, *rtracklayer: an R package for interfacing with genome browsers*. Bioinformatics, 2009. 25(14): p. 1841-2.

81. Lawrence, M., et al., Software for computing and annotating genomic ranges. PLoS Comput Biol, 2013. 9(8): p. e1003118.

82. Lock, A., et al., PomBase *2018: user-driven reimplementation of the fission yeast database provides rapid and intuitive access to diverse, interconnected information*. Nucleic Acids Res, 2019. 47(D1): p. D821-d827.

83. Hahne, F. and R. Ivanek, Visualizing Genomic Data Using Gviz and Bioconductor. Methods Mol Biol, 2016. 1418: p. 335-51.

84. Sætrom, P. and E.B. Stovner, epic2 efficiently finds diffuse domains in ChIP-seq data. 2019.

85. Canzio, D., et al., A conformational switch in HP1 releases auto-inhibition to drive heterochromatin assembly. Nature, 2013. 496(7445): p. 377-81.

86. Yamanaka, S., et al., RNAi triggered by specialized machinery silences developmental genes and retrotransposons. Nature, 2013. 493(7433): p. 557-560.

